# Integrins pattern the Drosophila embryonic neuroepithelium by influencing progenitor morphodynamics, division and position

**DOI:** 10.1101/2025.09.10.675109

**Authors:** Lamiya Dohadwala, Anupriya Garg, Maithreyi Narasimha

**Author notes:** Corresponding Author/Lead contact:.

## Abstract

Gene regulatory networks that confer cell fate and intracellular force generation mechanisms that drive cellular morphodynamics guide the patterning of tissues, but the nature of their interplay is poorly understood. We explore this during Drosophila embryonic neurogenesis whose first step, the delamination of a neuroblast from the surface epithelium, resembles an epithelial to mesenchymal transition (EMT). Using real-time, 3D confocal microscopy combined with quantitative morphometry, we identify robust morphometric state transitions that culminate in the basal delamination of the neuroblast (NB) from an equipotent epidermal proneural cluster, and find that mitotic rounding occurs concomitantly with delamination. We uncover marked heterogeneity and reduced collectivity in the morphodynamics of the NB and its nearest neighbours, and suggest their origins in the morphogenetic activity in the extending germband. We show that the appearance and distribution of the NB and GMC fate determinants Deadpan/dpn and Prospero/pros respectively correlate temporally with distinct morphometric states. We identify changes in cell-cell and cell-substrate adhesion that accompany neuroblast delamination and division and show that the adhesive microenvironment provided by integrin-ECM interactions patterns the neuroepithelium by influencing the spatiotemporal control of cell shape, division and position.

## Introduction

A large number of studies across the plant and animal kingdoms have established the reliance of cell fate specification during development on top-down control by gene regulatory networks (GRNs) including transcription factors and signaling pathways that define cell-fate specific transcriptomes (Arnone & Davidson, 1997; Davidson & Levine, 2008; Mircea & Semrau, 2021). The advent of quantitative high-resolution microscopy has enabled both the visualization of the tissue microenvironment in which cell fate specification occurs and the influence of fate determinants on cell and tissue morphodynamics (Chan et al., 2017; Copperman et al., 2023). An exemplary case in point is the specification and patterning of the Drosophila mesoderm which is accompanied by distinct changes in the transcriptomes and proteomes of mesodermal cells, and is regulated by the transcription factors Twist and Snail which serve as tissue selector genes (Calderon et al., 2022; Gomez et al., 2024; Leptin, 1991; Stern et al., 2022). Developmentally regulated expression of these transcription factors prefigures and instructs distinct aspects of cell shape changes accompanying the eventual internalization of the mesodermal germ layer by invagination (ventral furrow invagination/VFI; Martin et al., 2009; Schäfer et al., 2014; Sweeton et al., 1991; Weng & Wieschaus, 2016). An exogenous mechanical stimulus can compensate for the absence of endogenous deformation and induce the expression of Twist and Snail, suggesting that morphogenetic forces might feedback regulate transcription and cell signalling both in the native and perturbed states (Farge, 2003; Mitrossilis et al., 2017). There is also mounting evidence for the mechanical regulation of gene expression and through it, cell fate, and for the influence of morphogenetic activity and cell geometry on cell signalling (Brouzés & Farge, 2004; Chan et al., 2017; Falo-Sanjuan & Bray, 2022; Shaya et al., 2017). Whether and how morphogenetic activity at the cell and tissue scales influences fate determination, and what the nature of the interplay between gene expression, morphodynamics and the patterning of tissues is, remains poorly examined.

The epithelial to mesenchymal transition (EMT) is a cellular process accompanied by the loosening of contacts of an epithelial cell from its nearest neighbours, and the acquisition of a mesenchymal character that endows it with migratory/invasive potential (Thiery et al., 2009). It is widely deployed during development, homeostasis and disease, and depends on the EMT transcription factors that in turn induce changes in the expression and distribution of adhesion proteins, cell signalling molecules, and cytoskeletal components. Emerging evidence suggests that the expression of EMT transcription factors as well as the transition itself can be influenced by the mechanical microenvironment including but not limited to the mechanical forces generated by cell interactions with the extracellular matrix (Horta et al., 2023). The invagination of the VF described above can be thought of as a collective EMT, in that a band of cells held together by cell adhesion, albeit weak, collectively exhibit changes in cell shape, adhesion, apicobasal polarity and gene expression characteristic of an EMT and rely on the expression of the EMT transcription factors Twist and Snail (Leptin, 1991; Schäfer et al., 2014). The nature of the interplay between gene regulatory programs for fate specification and cell morphodynamics accompanying tissue morphogenesis however remains remarkably poorly understood except in a few contexts of *in vivo* EMT (Amack, 2021; Martin et al., 2009; Schäfer et al., 2014; Theveneau & Mayor, 2012; Weng & Wieschaus, 2016). That the microenvironment is an important determinant of differentiation outside of an EMT is borne out in the identification of stem cell niches that are typically thought to create chemical microenvironments that restrict cell signaling locally (Chacón-Martínez et al., 2018), and in the influence of the mechanical microenvironment including substrate stiffness on the regulation of cell fate (Engler et al., 2006; Smith et al., 2018). Whether and how the mechanical microenvironment contributes to cell fate specification *in vivo* and what the origin of such mechanical influences are during native/unperturbed development remains poorly explored.

One developmental context in which single cells loosen their contacts from their neighbours, move basally and acquire new fates as they do so is during embryonic neurogenesis in Drosophila. This cellular process, neuroblast (NB) delamination, generates the founders of the stereotypically patterned Drosophila nervous system. NB specification relies on contact and Notch signalling-dependent lateral inhibition that singles a NB out from each of the basic helix-loop-helix transcription factor achaete (ac) expressing proneural clusters in the surface ectoderm (neuroectoderm) in which the NB forms at the centre of a rosette while its nearest neighbours remain epidermoblasts (Hartenstein & Wodarz, 2013). The delamination of the NB bears resemblance to an EMT in being associated with both a cell morphodynamic state change and a cell fate change. This notion is also consistent with the recent identification of the EMT transcription factor of the Snail family, worniu, as the ‘missing’ proneural gene (Arefin et al., 2019). While the asymmetric division of the NB after delamination has received considerable attention (Barros et al., 2003; Campos-Ortega, 1997), remarkably little is known about the 3D morphodynamic changes that accompany the delamination of the NB other than its anisotropic apical loss (Simões et al., 2017). Also, whether or how the gene regulatory program for NB specification temporally correlates with or influences its delamination and the final pattern of the nervous system, and whether the morphodynamic changes feedback regulate gene expression has not been previously investigated in the context of early neurogenesis. Transcriptional regulation and the activity of enhancers in the embryonic mesectoderm, including those that are dependent on Notch signalling, have been recently shown to be sensitive to morphogenetic activity accompanying gastrulation (Falo-Sanjuan & Bray, 2021). NB delamination and differentiation in the Drosophila embryo occur in the ectoderm during the morphogenetic movements of germband extension and retraction when widespread changes in tissue tension and cell rearrangements in the plane of the epithelium also occur. Whether and how these movements influence NB fate specification also remains unclear.

Single cell delamination can also occur in cells that have already become committed to a particular fate, where it has been demonstrated to contribute to force generation, the maintenance of tissue tension, cell number homeostasis and cell quality control (Eisenhoffer et al., 2012; Marinari et al., 2012; Toyama et al., 2008). Earlier work from our lab has demonstrated that delamination of single cells in the amnioserosa during dorsal closure is accomplished by changes in cell shape, cell adhesion and cytoskeletal organization, and that its number and position are influenced by both intracellular and exogenous mechanical forces and chemical signals (Guru et al., 2022; Meghana et al., 2011; Muliyil et al., 2011; Muliyil & Narasimha, 2014; Saravanan et al., 2013). Whether the morphodynamic features of cell delamination differ depending on whether or not it is associated with a transcription factor dependent fate change, and conversely whether all delaminations involve an EMT-like transcriptional program remain unresolved.

Integrin-ECM interactions are crucial regulators of morphodynamics at both cellular and tissue scales. In addition to providing a structurally supportive substrate for cells and cell sheets, these interactions generate and transduce mechanical and chemical signals and enable communication between cells and their external environment (Wu et al., 2023). The diversity of integrin-ECM interactions, generated by the differential deployment of ECM ligands, integrin heterodimers, and adapter proteins, enable the regulation of subcellular processes including the modulation of substrate composition and stiffness, the generation of intracellular and traction forces through cytoskeletal remodelling, the regulation of cell adhesion and polarity, and the control of cell signalling and gene expression (Engler et al., 2006; Green et al., 2018; Kechagia et al., 2019; Klapholz et al., 2015; Winograd-Katz et al., 2014). These subcellular functions in turn influence cell proliferation, survival, morphodynamics and fate. It has become increasingly evident that these interactions contribute character to the ‘microenvironment’ (Wu et al., 2023). The transcriptionally regulated cell-cell adhesion switch from E– to N-Cadherin is a hallmark of an EMT that is believed to loosen cells (Thiery et al., 2009). In flies and mice, alterations in E-Cadherin levels can alter the numbers of cells allocated to distinct fates, but whether changes in cadherin type or levels influence the acquisition of specific fates remains unclear (Schäfer et al., 2014; Stephenson et al., 2010).

In this work, we use a combination of real-time confocal microscopy and quantitative morphometry to uncover the spatiotemporal hierarchies in the link between cell morphodynamics, cell fate acquisition and cell position in the early patterning of the Drosophila embryonic neuroepithelium. We uncover the dynamic distribution patterns of cell-substrate interactions and their role as placeholders and timekeepers during early neurogenesis.

## Results

### A robust sequence of morphometric transitions accompanies embryonic neuroblast specification

The founders of the stereotypically patterned embryonic nervous system in Drosophila, the embryonic neuroblasts (NB), have their origins in the proneural competence fields (marked by the achaete/ac expressing cell clusters) in the embryonic ectoderm from which they are singled out for delamination through Notch dependent lateral inhibition (Fig. 1C). Their specification occurs in five waves over three hours spanning from stage 8 to stage 11 of embryogenesis (Artavanis-Tsakonas et al., 1999; Hartenstein & Campos-Ortega, 1984; Hartenstein & Wodarz, 2013). The basal positioning of the neuroblast which retains achaete expression relies on its delamination from the surface ectoderm (dorsal ectoderm at germband extension and retraction in this work; Fig.1 C, D), and is followed by its asymmetric division into a larger NB and a smaller GMC (Knoblich, 2008). Early NBs and their derivatives occupy more basal positions in the progressively dorsoventrally layered neuroepithelium in a manner reminiscent of the dorsal-ventral patterning in vertebrates (Isshiki et al., 2001). The neuroblasts and their derivatives express transcription factors including Deadpan/dpn (Fig 1 D, E; Bier et al., 1992) that confer fate/identity, and Hunchback/hb, Castor/Cas, Kruppel/Kr, and Grainyhead/Grh that indicate their birth order (Isshiki et al., 2001).

**Figure 1:**
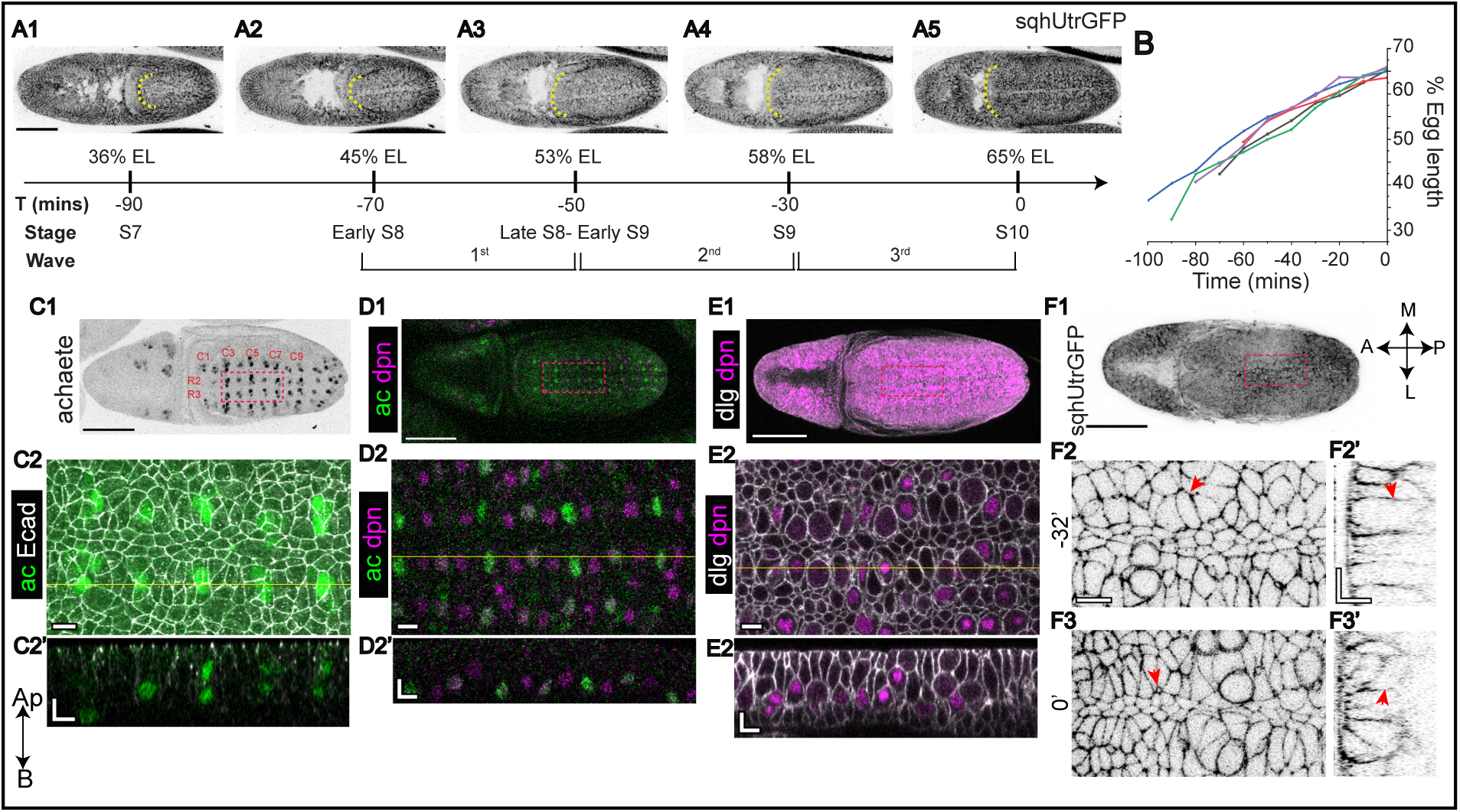
Experimental paradigm for the analysis of embryonic neurogenesis. (A1-A5) Chronology and dynamics of germband extension showing germband length represented as percentage of egg length (EL) in stills from real-time confocal microscopy of the dorsal ectoderm in a representative embryo expressing SquashUtrophin::GFP. Dashed lines in A mark the extending anterior boundary of the germband, and the embryonic stage and time windows within which the waves of neurogenesis proceed are shown below it. Waves 2-3 of neurogenesis were used for all morphometric analyses. (B) Dynamics of fractional germband length changes measured from real-time movies of five SquashUtrophin::GFP expressing embryos and plotted in a retrospective time scale (t0 marks the fully extended germband). (C) A *w^1118^* embryo immunostained for achaete (ac) (black in C1, green in C2) to mark the proneural clusters arranged in stereotypical rows (R) and columns numbered in red. C3-C7 and R2-R3 (marked on C1) were used as the ROI for all analyses. (C2) High magnification views of the above mentioned ROI of an embryo stained with achaete (green) and E-Cadherin (ECadh, grey) along with an orthogonal section (C2’) through the dotted line. (D1) A SquashUtrophin::GFP embryo stained for achaete/ac (green) and deadpan/dpn (magenta) with an inset magnified in D2, along with its orthogonal section through the dotted line (D2’). (E1) A *w^1118^* embryo immunostained for dlg (grey) and dpn (magenta). The inset in E1 is magnified in E2 to show heterogeneities in dpn expression. An orthogonal section through the dotted line in E2 is shown in E2’. ( F1) A low magnification image of a SquashUtrophin::GFP embryo. (F2, F3) High magnification views of the inset in F1at two time points approximately 30 mins apart. Red arrowheads mark a delaminating NB. Scale bars: 100 μm (A1-A5, C1-F1); 10 μm (C2-F3’). In the axis labels, Ap-apical, B-Basal, A-anterior, P-posterior, M-medial and L-lateral.

We examined the morphodynamic changes accompanying neuroblast delamination in dorsal ectodermal proneural clusters (neuroectodermal fields in the future caudal germband) during the second and third waves of neurogenesis (end GBE and early GBR stages, germband occupying 58-65% of the embryonic length/EL; Fig. 1 A, B) in real-time using SquashUtrophin::GFP embryos in which the decoration of cortical actin facilitated the visualisation of cell shapes in both XY and YZ. NBs from these clusters were identified on the basis of their complete detachment and basal delamination from the dorsal ectoderm and their subsequent asymmetric division, and tracked retrospectively (Fig. 1 F, see methods).

Prior to the onset of apical constriction, NBs exhibit a columnar morphology with a nearly uniform cross-sectional area along the apicobasal axis, and are indistinguishable from their epidermal neighbours. We call this state S1 (Fig. 2 A1’-A3’’, B1’-B4’’, C). While the duration or start of the S1 state cannot be determined, apical expansion was observed in the 30-60 minutes prior to the onset of apical constriction in a large fraction of epidermal cells that subsequently delaminated basally (neuroblasts/NB; 63%, n=28 cells from 5 embryos; compare Fig. 2 A1’-A3’ and Fig. 2 B1’-B3’). The prior occurrence of apical expansion did not however alter the subsequent morphometric changes accompanying delamination. The onset of apical constriction and the expansion of the basolateral region of the cell, characterise the frustum-shaped S2 state in which the width at the NB base is approximately twice that at its apical end (Fig. 2 A4-A5, B5, C). Constriction initially observed only at the apical end in S2 subsequently extended laterally leading to the S3 state which is characterised by a prominent, narrow, elongated stalk that is clearly distinct from the bulbous base (Fig. 2 A6-A7, B6-B7, C). In the early S4 state, the NB is largely bulbous with a very short, pinched, residual stalk that ultimately diminishes to form a spherical NB that lies beneath the dorsal ectoderm from which it originated (late S4; Fig. 2 A8-A10, B8-B10, C). From the onset of apical constriction, delamination (S2 to S4 states) takes approximately 45 minutes. The subsequent state S5, identified by the appearance of a small basal bud (Fig. 2 A11, B11), is followed by the completion of division and the formation of two daughter cells: a larger NB, which marks the S6 state, and a smaller ganglion mother cell (GMC) (Fig. 2 A12, B12, C). Division is completed within five minutes of delamination.

**Figure 2:**
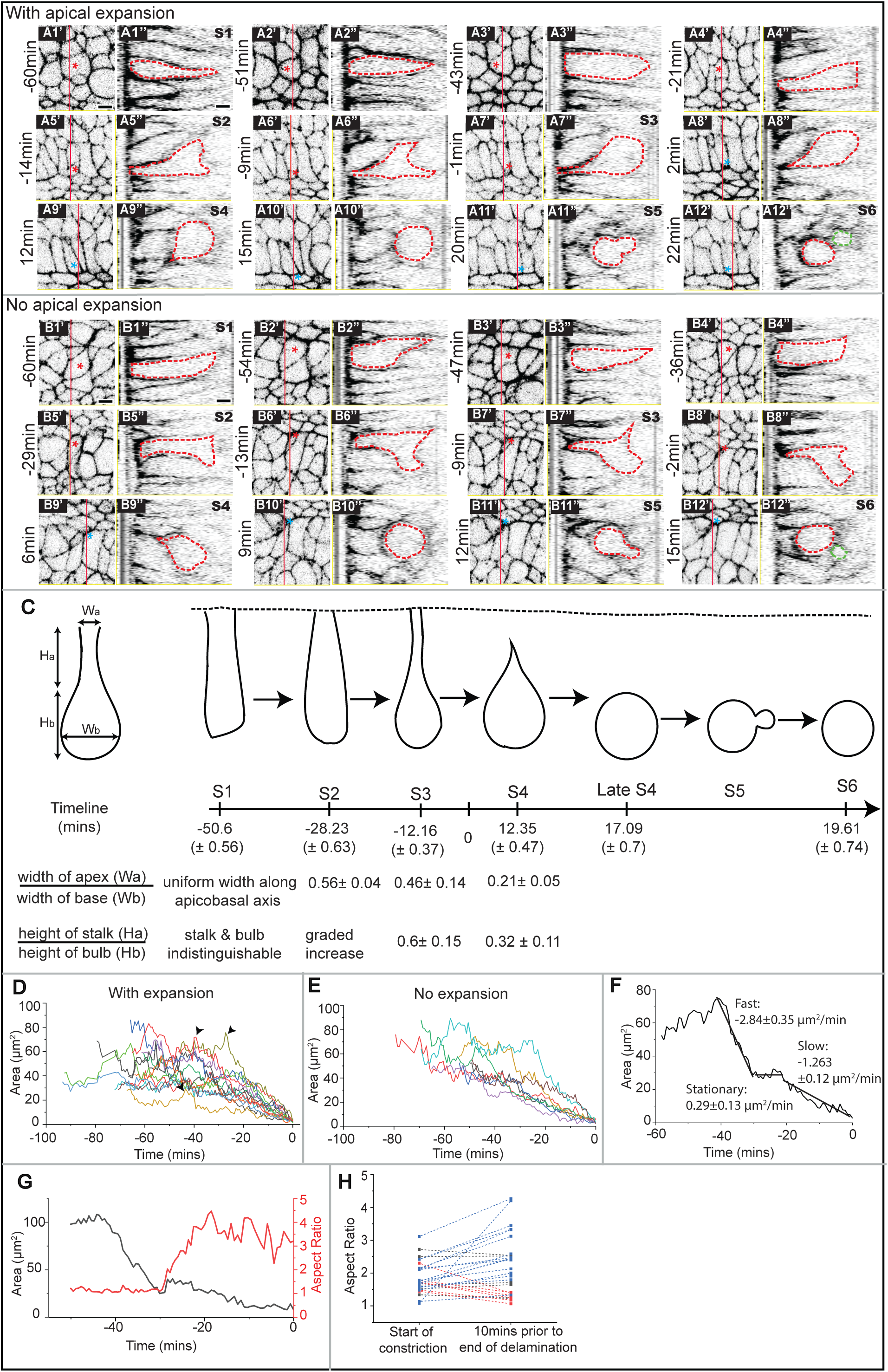
Morphodynamics accompanying neuroblast delamination. (A, B) Time-lapse images of maximum intensity projections of the apical slices of delaminating neuroblasts and their orthogonal sections in wild-type embryos expressing SquashUtrophin::GFP showing cell shape changes accompanying NB delamination. Two representative NBs are shown, one with (A, A1’-A12’’) and one without (B, B1’-B12’’) apical expansion before constriction. t0 corresponds to the loss of apical cell membrane (apical area<3 μm²). Red dashed lines mark cell boundaries of delaminating neuroblasts in the orthogonal views. Red asterisks indicate NBs of interest while blue asterisks indicate the position of the NB after it has lost its apical membrane. Green dashed line in A12’’, B12’’ indicates the GMC formed after asymmetric division. Apical is on the left in all images. Scale bar: 10 μm. (C) Morphometric features that characterise the six states accompanying NB delamination. A time line depicting the chronology of delamination and the ratios of the width and height of the stalk and bulb that characterise these states is also shown (n=10 cells from 3 embryos; data presented are means ± SEM). (D, E) Apical area dynamics of delaminating NBs with expansion prior to net constriction (D, arrowheads indicate the expansion phase, n=14 cells from 5 embryos) or without expansion (E, n=8 cells from 4 embryos). (F) Representative plot of apical area dynamics of NB showing fast, stationary and slow phases of constriction. Rates shown alongside are mean ± SEM (n=12 cells from 4 embryos). (G) Apical area and aspect ratio changes accompanying delamination in a representative NB. t0 depicts the first instance when the apical area of NB falls below 3 μm². (H) Aspect ratio of delaminating NBs at the start of constriction and 10 mins prior to end of delamination. Red, blue and black line show respectively decreasing, increasing or unchanging aspect ratios.

Thus, the ‘specification’ of the NB is accompanied by a continuum of cell shape changes with clearly identifiable transitional states occurring over a period of approximately 60 minutes (Fig. 2C). Shape changes leading up to delamination occur comparatively slowly, and asymmetric cell division follows rapidly thereafter. These morphodynamic changes suggest that mitotic rounding may be initiated prior to the culmination of delamination, a validation of which require an assessment of cell volume and cell cycle marker dynamics accompanying the morphometric changes.

### Apical constriction in delaminating neuroblasts is anisotropic and heterogeneous

Quantitative morphodynamic analysis of a number of NBs revealed heterogeneity in apical constriction patterns. In half of the NBs analyzed (53%, n=15), three phases could be discerned: an initial fast phase (areal reduction of ∼3 μm²/min over approximately 10 minutes), a brief stationary phase (5 minutes) with little or no net area reduction, and a longer (25 minutes) slow phase where area reduction rates were two fold lower (∼1.3 μm²/min, Fig. 2 F, S1 A1-A15). Constriction in the remaining 47% of NBs proceeded in a single phase at the slower rate of constriction (1.22+/− 0.56 μm²/min, n=13; Fig. S1 A16-A28). The apical shapes of the majority of NBs examined became progressively more anisotropic, consistent with earlier reports (Simoes et al, 2017). Starting with an average aspect ratio of 1.8 at the onset of apical constriction, approximately half of the NBs analysed (57%, n=16, Fig. 2 G, H, 3 A-C, S2 A13-28) became increasingly anisotropic (mean aspect ratio of 2.7) ten minutes prior to the end of delamination. Of the remainder, half (21%, n=6, Fig. 2H, S2 A7-12) did not show a further increase, and half (21%, n=6) modestly reduced the anisotropy (mean aspect ratio of 1.26, Fig. 2H, S2 A1-A6). These observations reveal that apical constriction accompanying NB delamination is heterogeneous and suggest that morphogenetic activity in the extending germband may contribute to the observed differences.

### Rosettes around delaminating neuroblasts are radially asymmetric and exhibit neighbour losses and gains

During cell delamination or extrusion, in other contexts, the nearest neighbours (NN) collectively change their shapes and alignment as the delaminating cell constricts, to form a nearly radially symmetrically patterned rosette with the delaminating cell at its centre. The NNs contribute essential forces for delamination (Meghana et al., 2011; Rosenblatt et al., 2001; Saravanan et al., 2013; Toyama et al., 2008). An examination of the morphodynamics in the NNs of delaminating NBs revealed marked heterogeneity and differed from delamination elsewhere in three ways. First, the rosettes formed around a delaminating NB are asymmetric and lack a clear radial axis at their apical ends owing to the heterogeneity in the apical shapes and sizes of its NNs (Fig. 3 A-C, S3). Second, the number of nearest neighbours (at the apical end) and therefore, the sidedness of the NBs changes during the course of delamination. On average, neighbour loss occurs more frequently than neighbour gain, evident from the observation that in the majority of cohorts analysed, the number of neighbours towards the end of delamination is lower than at the onset of constriction (Fig. 3 A1-A7, 3E). This neighbour loss occurs more frequently in the 20 minutes prior to delamination, and can occur more than once in a NB cohort (Fig. 3F). Cell rearrangements leading to neighbour exchange account for neighbour loss in the majority of cases (92.8% of cohorts, 73/171 NNs, Fig. 3G). In a smaller proportion (42.8% of cohorts, 15/171 NNs, Fig. 3G), a delaminating neighbour reduces the neighbour number (Fig 3A1-A7, B1-B7, H). In a very small fraction of NB cohorts (7%; 4/56 NNs), a gain of neighbours accounted for the increase in sidedness of the NB whose frequency was also highest in the 20 mins prior to delamination (Fig. 3 F, H). Neighbour gains were found to result from cell rearrangements (50% of cohorts, 16/171 NNs), or cell division leading to the formation of new contacts with both daughter cells (Fig.3 C1-C7). Third, NB delamination was also accompanied by apical expansion in a third of its nearest neighbours (1.8 cells in a cohort of an average of 6 cells, from a total of 83 NN from 28 cohorts). On average, the apical area increased 1.2 fold by the end of delamination. Neighbour loss was often correlated with the expansion of an adjacent NN (Fig. 3A1-A7, D). These observations underscore the morphodynamically heterogenous neighbourhood within which NB delamination occurs and raise the possibility that morphodynamics may influence fate specification autonomously or non-autonomously. Additionally, they identify delaminations occurring in neighbouring cells.

**Figure 3:**
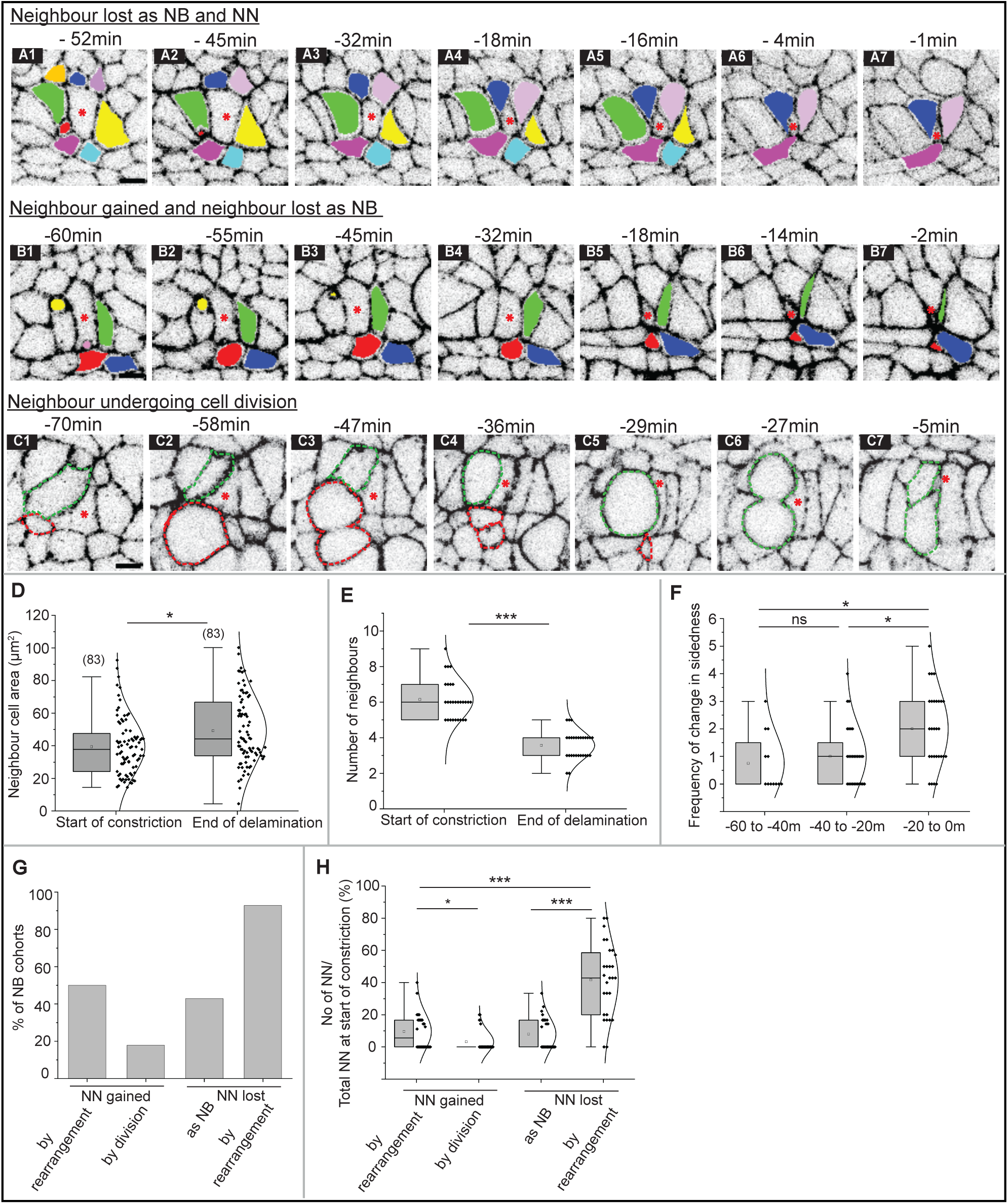
Nearest neighbour morphodynamics accompanying neuroblast delamination. (A-C) Stills from real-time confocal microscopy of three delaminating neuroblast cohorts in the dorsal ectoderm in embryos expressing SquashUtrophin::GFP showing morphodynamic heterogeneities in the NB neighbourhood. t0 corresponds to the loss of apical cell membrane (<3 μm^2^). Red asterisks and colour-filled cells indicate, respectively, the delaminating neuroblast and its neighbours. (A1-A7) Loss of neighbours as a NB (red) or by cell rearrangements (NN; green, yellow, cyan). (B1-B7) NN gained (blue) and neighbours lost as NB (yellow, red, green). (C1-C7) Neighbours undergoing cell division (red and green dashed lines). Scale bars: 10 μm. (D) Nearest neighbour apical cell area and (E) number at the onset of net apical area constriction and the end of delamination of the NB in each cohort (n=28). (F) Number of times neuroblasts changed their sidedness during the course of delamination. t0 corresponds to the loss of apical cell membrane (<3 μm^2^). (G) Percentage of NB cohorts showing neighbours lost or gained by cell rearrangements, division or delamination. (H) Nearest neighbours gained or lost by rearrangement, delamination or division expressed as a fraction of the total population of neighbours at the start of net constriction in each NB cohort. Twenty-eight NB cohorts from 5 embryos were considered for analysis. In the box plots in D-F, H boxes show median (horizontal line) ± interquartile range; the mean is indicated by ‘□’ and the sample size is in brackets. The Mann–Whitney test was used for statistical analysis (*-p < 0.05, **-p < 0.01, ***-p < 0.001, and ns-not significant).

### The subcellular distribution of NB and GMC fate determinants Deadpan and Prospero show temporal correlations with the morphometric state of the NB

We chose to examine the temporal patterns of expression of two fate determinants in the context of embryonic neurogenesis. Deadpan (dpn), a bHLH transcriptional repressor, and a member of the Hairy and Enhancer-of-Split (Hes) family, is expressed specifically in neuroblasts and is involved in maintaining NB self-renewal (Bier et al., 1992). Prospero (pros), a member of the homeobox family exhibits dual localization. Initially localized in the basal cortex of NBs, it becomes nuclear in the GMCs resulting from its asymmetric division. The cortical asymmetry in the NBs depends on the stage of the cell cycle and on myosin activity (Barros et al., 2003; Mora et al., 2018) and enables the unambiguous identification of both populations.

We monitored the appearance of dpn protein in the dorsal ectoderm during stages 9-10 of embryogenesis using an antibody against it (Fig. 1E). Deadpan protein was undetectable in cells in the ectodermal surface layer. It was first detected in cells in the early S4 morphometric state where it colocalized with DAPI (Fig. 4A, A1, A1”, A1”, B). A small population of S4 NBs did not show dpn expression (Fig. 4C) suggesting that dpn expression begins in the S4 state. We identified heterogeneities in dpn distribution and intensity that correlated with the morphometric state. In the majority of S4 state NBs, dpn was localized to the large nucleus of the NB that occupied a large fraction of its cross-sectional area barring a slim rim of cytoplasm (∼70%, n=82; Fig. 4 A1, A1’, A1’’, 4B, 4C). In the S5 state, the dpn signal was predominantly (∼75%, n=41) faint, diffuse and cytoplasmic, and the DNA was organized into mitotic figures (Fig. 4A, A2, A2’, A2’’, 4C). This strongly suggests that the diffuse cytoplasmic dpn might be associated with nuclear envelope breakdown prior to the onset of asymmetric cell division. Cells exhibiting compact, intense nuclear dpn were also found, that we believe mark the self-renewed NB daughter (S6) following the asymmetric cell division (Fig. 4A, A3, A3’, A3”, 4C). This notion is strengthened by the observation that in some S5 cells with the basal bud (Fig. 4C), compact nuclear dpn is seen in the larger of the two daughter cells of the asymmetric division in which karyokinesis but not cytokinesis is complete. The S6 morphometric state was heterogeneous with respect to dpn distribution. One class of cells exhibited an intense nuclear localization of dpn as described above, while the other showed the enlarged nuclear dpn described for newly delaminated NBs in the S4 state (Fig. 4C). The latter could, in fact, be newly delaminated NBs that have lost their apical stalk (late S4) and show a rounded S6-like morphology (Fig. 4C). These findings indicate that dpn protein is undetectable in NBs until they have moved basally, and suggest that weakened adhesive contacts with neighbours or successful detachment/delamination may be a necessary trigger for deadpan expression. Our observations do not rule out the possibility that dpn transcription is initiated earlier.

**Figure 4:**
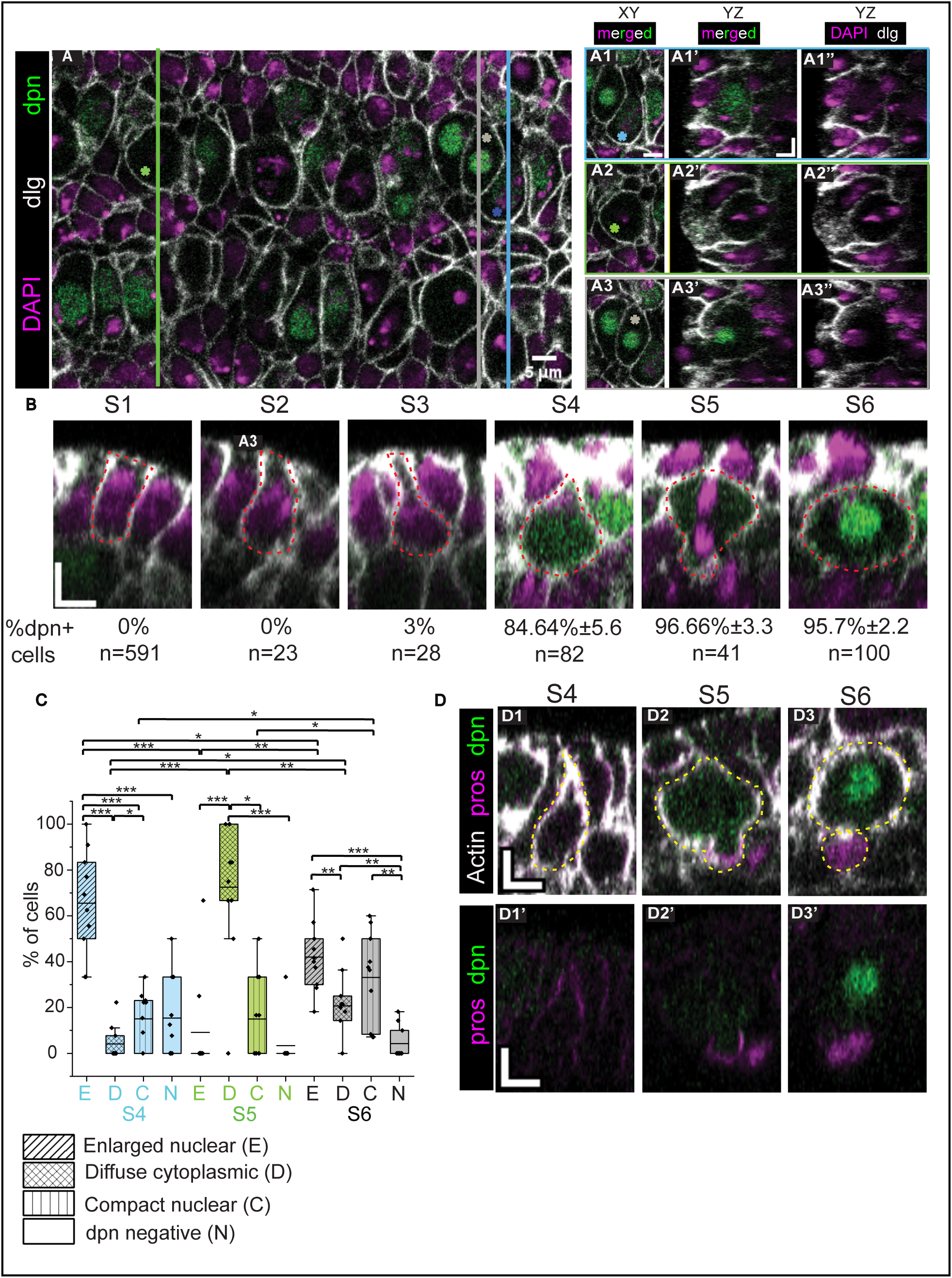
Heterogeneity in Deadpan expression correlates with the morphometric state of the neuroblast. (A) A Single slice at 15 μm depth from the dorsal vitelline membrane of a stage 10 *w^1118^* embryo stained with dlg (grey), dpn (green) and DAPI (magenta). The coloured vertical lines (blue, green and grey) mark the sites of the orthogonal (YZ) sections in A1 to A3, that cut across 3 cells (marked by the blue, green and grey asterisks) to showcase heterogeneities observed in Dpn expression that are also associated with different morphometric states. YZ orthogonal views of cells showing enlarged nuclear Dpn (A1-A1’), diffuse cytoplasmic Dpn (A2-A2’), and compact nuclear Dpn (A3-A3’) are shown, apical is to the left in all images. (B) Temporal correlation of Dpn distribution patterns in delaminating NBs in the S1-S6 morphometric states seen in the YZ orthogonal views of NBs (dashed red line marks the outline of the cell of interest; apical is on the top in all images). Their relative frequencies of occurrence in each morphometric state (percentage of Dpn+ cells of the total number of cells in that state) is provided below the images (n= 10 embryos). Scale bars: 5 μm. (C) Quantitative correlation between the morphometric states and Dpn distribution heterogeneity depicted as percentage of cells showing a given Dpn distribution pattern or no Dpn expression within the S4 (blue), S5 (green) and S6 (grey) morphometric states. In the box plots in C, boxes show mean (horizontal line) ± interquartile range. The Mann–Whitney test was used for statistical analysis (*-p < 0.05, **-p < 0.01, ***-p < 0.001, and ns-not significant). (D) Orthogonal (YZ) sections w118 embryos stained for Pros (magenta), Dpn (green) and actin (grey) to show the temporal correlation of Prospero distribution patterns in delaminating NBs in the S4-S6 morphometric states (dashed yellow lines mark the outline of the NB and the GMC; apical is on the top in all images). Scale: 5μm.

**Figure 5:**
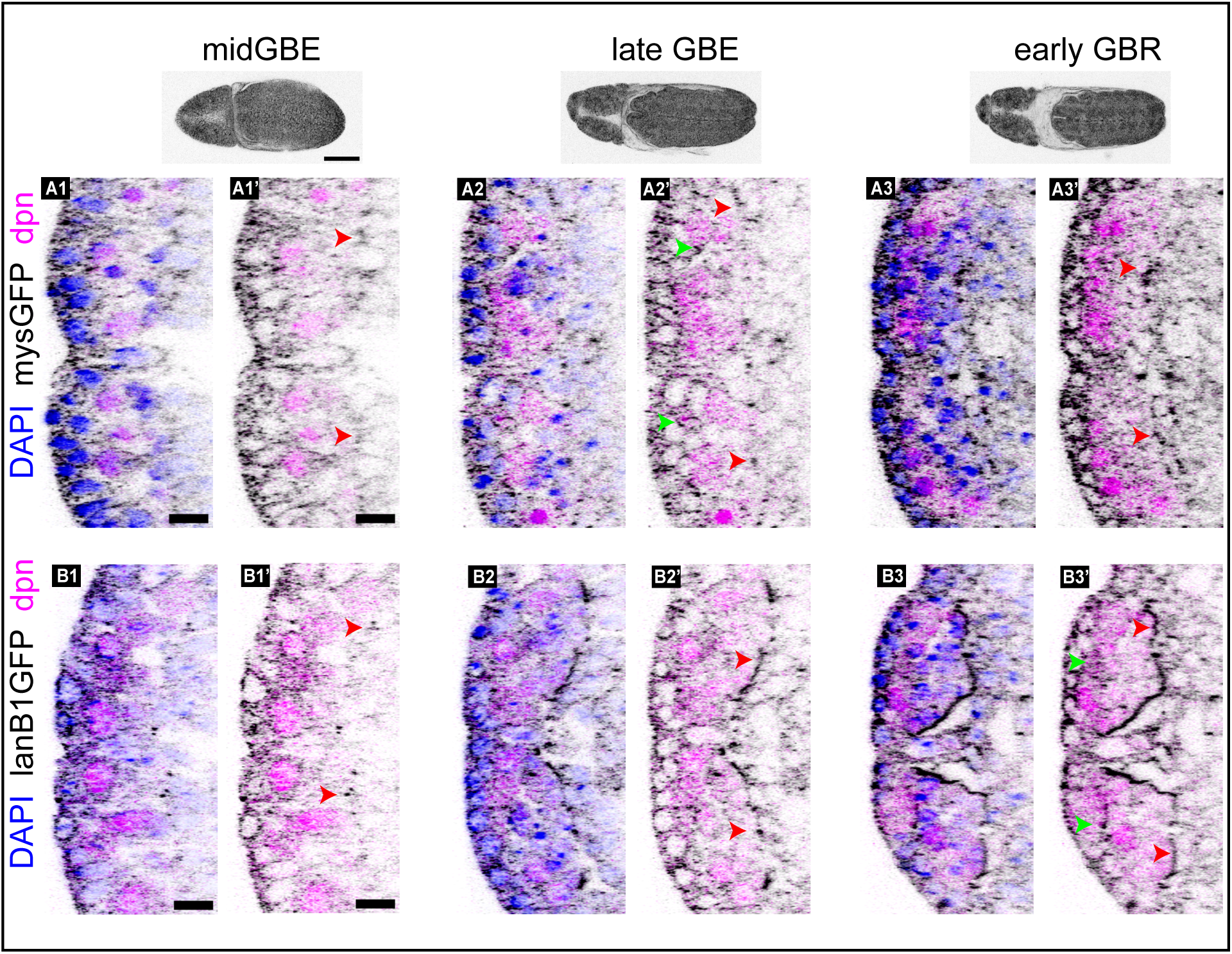
Temporal regulation of integrin and laminin distribution during early embryonic neurogenesis. Orthogonal sections of mysGFP (A, grey) or lanB1GFP (B, grey) embryos from mid GBE (A1, B1), late GBE (A2, B2) and early GBR (A3, B3) co-stained for Dpn (magenta) and DAPI (blue). Red and green arrowheads arrowheads show respectively enrichment of integrin and laminin at the neuroepithelium-mesoderm boundary or elsewhere within the neuroepithelium. In each image, apical is on the left. Scale: 10 μm. The stages of embryogenesis from which the orthogonal sections were obtained is shown in the low magnification images of embryos at the top. Scale: 100 μm.

We also monitored the expression of Prospero. As mentioned above, Prospero has been shown to act as a fate determining principle through its asymmetric basal cortical localisation in the NB, and its subsequent inheritance and nuclear translocation in the basal daughter that forms the GMC. We found a strong apical cortical enrichment of pros as has also been described previously (Spana & Doe, 1995), which correlated with the S4 morphometric state and preceded both its basal cortical enrichment (which was pronounced in the S5 state) and the appearance of nuclear dpn. This was followed by the appearance of nuclear pros in the GMC in the S6 state (Fig. 4D). The correlation of cortical pros distribution with distinct morphometric states (S4 to S6) raises the possibility that Prospero translocation from the apical to basal cortex might influence the acquisition of distinct cell fates.

### The emergence of a dorsoventrally layered neuroepithelium from single cells is accompanied by spatiotemporal changes in the distribution of integrin and ECM proteins

In addition to the morphometric states that are defined by cell shapes, embryonic neurogenesis is also accompanied by changes in the organisation of the neuroepithelium that forms beneath the dorsal ectoderm. This is characterised by changes in the position of the delaminated NB and its derivatives along the dorsoventral (DV) axis. The stereotypical DV patterning of the Drosophila embryonic neuroepithelium evident at embryonic stage 12 bears semblance to the layered mammalian cortex. Early-born NB derivatives occupy deeper (more basal) positions while later-born NB progeny localize more superficially (closer to the dorsal ectoderm) (Isshiki et al., 2001). Deadpan positive NBs are positioned just beneath the dorsal ectoderm, while GMCs and their progeny are positioned further basally. The GMC layer comprises a single row of cells at stage 9, expanding to 3-4 rows by stage 12 as NB divisions progress (Fig.7 A1, A2, C1, C2). By embryonic stage 12, the wild-type neuroepithelium comprises four to five rows of cells along the dorsoventral axis, is approximately 25 μm thick, and occupies the space between the dorsal ectoderm and the underlying mesoderm from which they could be distinguished by Prospero staining (which labels all GMCs and their neuronal derivatives occupying the deepest three neuroepithelial rows; Fig. 6 A3, A4). This progressive stratification is also evident in the location of hunchback/hb positive cells within the neuroepithelium. Hunchback which marks the first born NBs was expressed in newly delaminated dpn^+^ NBs in the basal layer in early stages (Stage 9-10; Fig. 7 E1-E4) but became restricted to deeper layers by stage 13 (mid GBR), when its expression no longer overlapped with dpn, indicating terminally differentiated neurons or GMCs, as previously described (Fig. 7 G1-G4). Remarkably, the neuroepithelial thickness remained relatively constant over this period, and the increase in the number of cells was accompanied by compaction, evident from the reduced spaces between individual cells.

**Figure 6:**
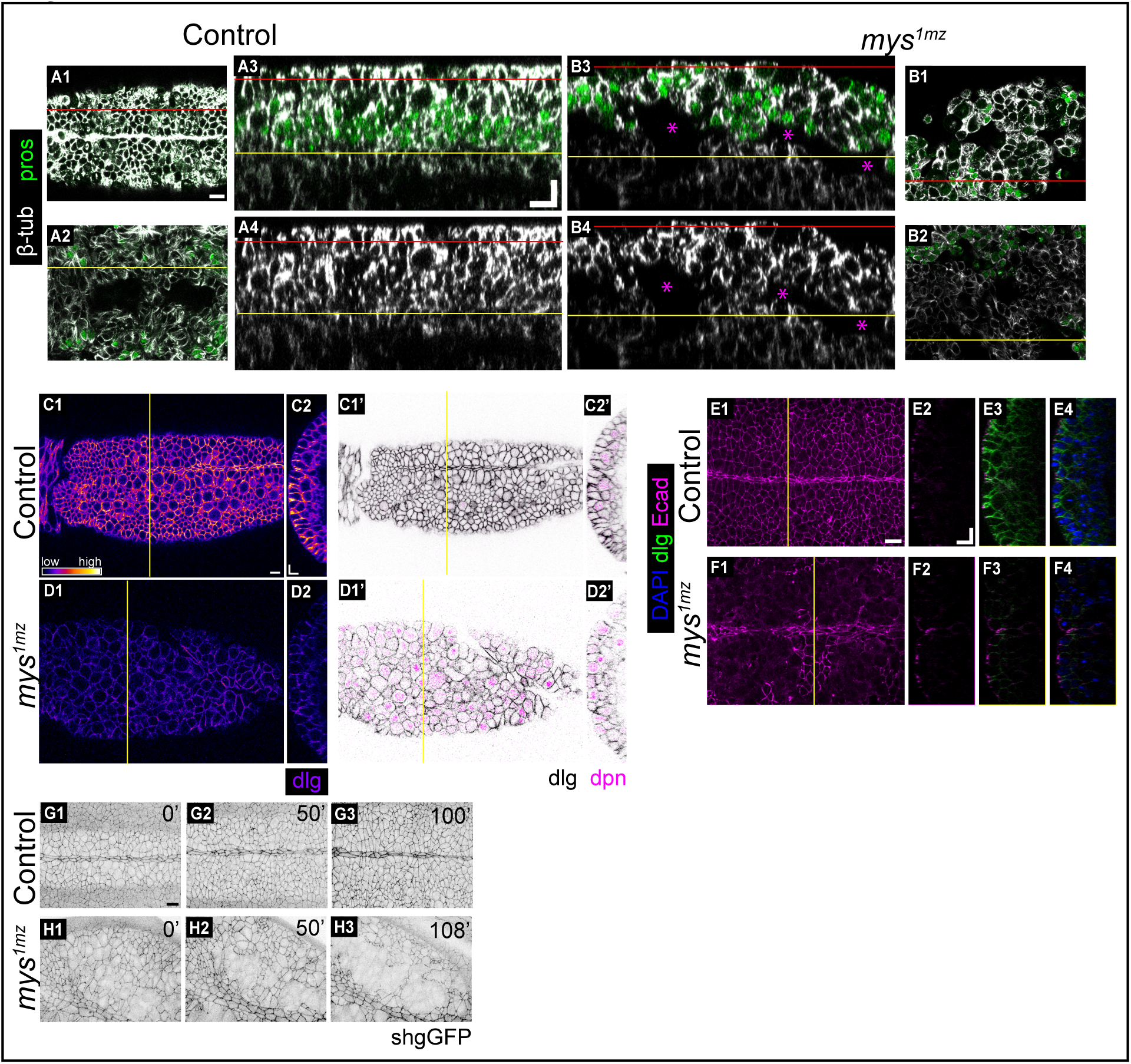
Defects in cellular organisation and stratification in the neuroepithelium in integrin mutants The dorsal neuroepithelium in heat-shocked control. (A) and germline clones of *mys^1^* (B) stained for β-tubulin (grey) and Prospero (green). Single slices at the level of the dorsal ectoderm (A1, B1) and mesodermal layer (A2, B2) in control (A1-A2) and *mys^1mz^* mutant (B1-B2) embryos and the orthogonal sections through the red lines running along the anterior-posterior axis (A3, A4, B3, B4). Differences in the apposition of the neuroepithelium and mesoderm, cell packing and tissue organization between controls (A3, A4) and *mys^1mz^* mutant (B3, B4) embryos are seen. Depth of the single slices and position of the orthogonal sections are marked by the red and yellow lines. Magenta asterisks indicate separation between the neuroepithelium and mesoderm. Anterior is to the left A1, A2, B1, B2 and apical is at the top in A3, A4, B3, B4. (C, D) Control (C) and *mys^1mz^* embryos (D) stained for dlg (intensity coded, fire in C1, D1 and grey in C1’, D1’) and Dpn (magenta in C1’, D1’) to reveal decreased intensity of dlg and the presence of dpn in the dorsal ectoderm in *mys^1mz^* (D1, D2, D1’, D2’) embryos compared to controls (C1, C2, C1’, C2’). C2, D2, C2’, D2’ are YZ orthogonal slices through the yellow line. Apical is to the left. (E, F) Control (E) and *mys^1mz^* embryos (F) stained for E-Cadherin (magenta), dlg (green) and DAPI (blue) and to reveal the patchy loss of E-Cadherin (magenta) in *mys^1mz^* mutants (F1-F4) compared to controls (E1-E4). E1, F1 are apical XY projections and E2-E4, F2-F4 are YZ orthogonal sections of the tissue cutting through the yellow lines in E1 and F1. Apical is to the left. (G, H) Time-lapse images showing apical projections of the dorsal ectoderm in control (G1-G3) and *mys^1mz^* mutant (H1-H3) embryos carrying the shotgunGFP transgene to show the progressive loss of ectodermal E-Cadherin in *mys^1mz^* embryos. In all images anterior is to the left. Scale: 10 μm.

**Figure 7:**
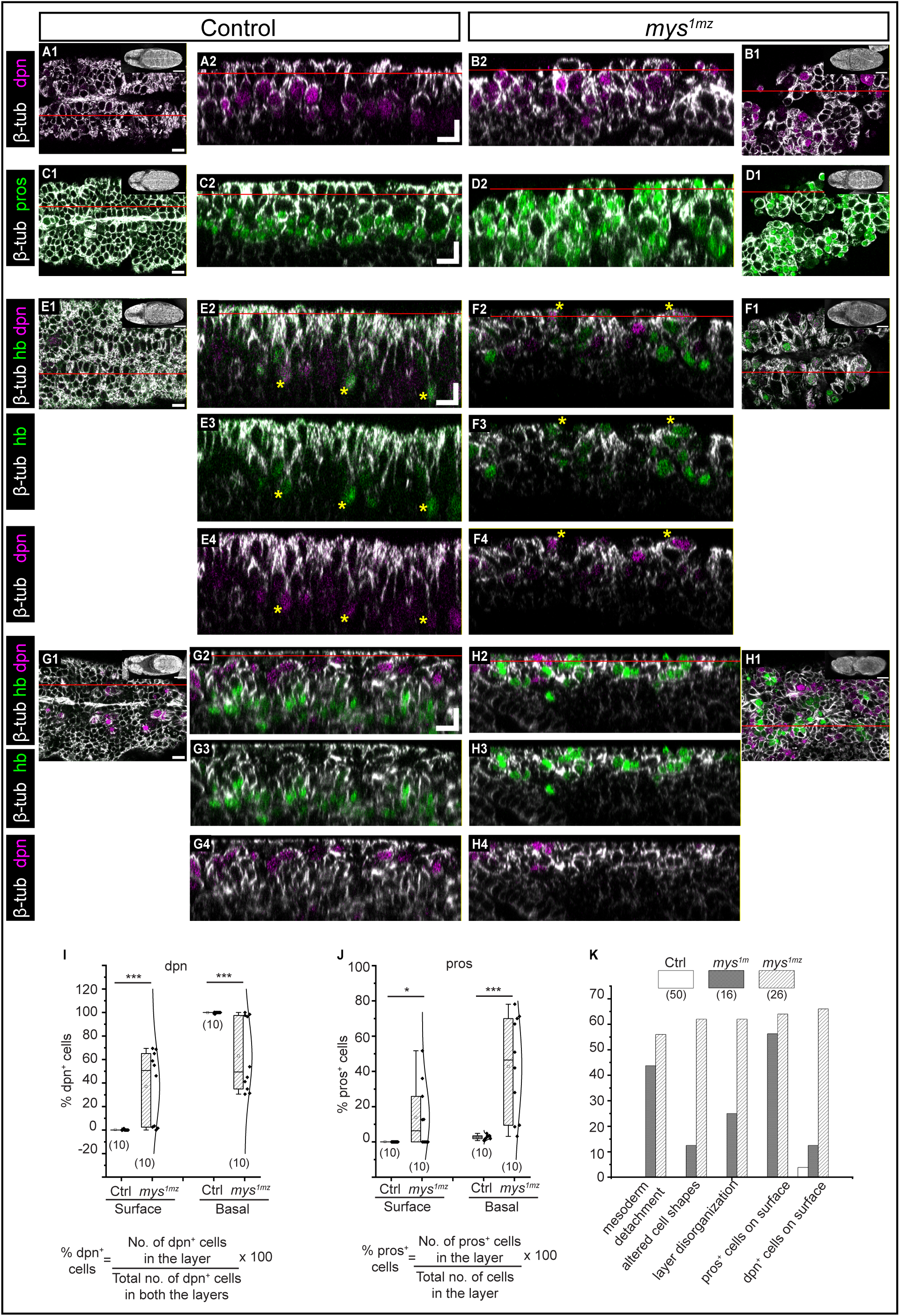
Defects in dorsoventral layering and neuroblast position in the neuroepithelium of integrin mutants Control. (A, C, E, G) and *mys^1mz^* mutant (B, D, F, H) embryos immunostained for β-tubulin (grey) and dpn (magenta) (A, B) or β-tubulin (grey) and prospero (green) (C, D) to reveal layer disorganisation and NB and GMC mispositioning in *mys^1mz^*mutant embryos (compare A, C with B, D). A2, B2, C2 and D2 are orthogonal sections through the red lines in A1, B1, C1 and D1(insets in A1-F1 are low magnification images showing embryonic stage). Control (E1-E4 and G1-G4) and *mys^1mz^*mutant (F1-F4, H1-H4) embryos at the end of GBE (E, F) or midway through GBR (G, H) stained with β-tubulin (grey), hunchback (green) and dpn (magenta) to reveal the disorganised layering in the neuroepithelium of *mys^1mz^*mutant embryos (F1-F4, H1-H4) compared to controls (E1-E4 and G1-G4). Red lines in E1, F1, G1 and H1 indicate the plane of the orthogonal sections in E2-E4, F2-F4, G2-G4 and H2-H4. Red lines in E2, F2, G2 and H2 indicate the XY plane of the sections in E1, F1, G1 and H1. Yellow stars indicate co-expression of hb and dpn. Anterior is to the left in XY views and apical is at the top in orthogonal sections. Scale: 10 μm. (I, J, K) Quantitative analysis of the percentage of dpn^+^ (I) and pros^+^ (J) cells on the surface or basally in controls and *mys^1mz^*mutants. In the box plots in I-K, boxes show median (horizontal line) ± interquartile range; the mean is indicated by ‘□’ and the sample size is in brackets. Mann– Whitney test was used for statistical analysis (*-p < 0.05, **-p < 0.01, ***-p < 0.001, and ns-not significant). K) Prevalence (relative frequency) of tissue and cell scale phenotypes in control, *mys^1mz^* and *mys^1m^* mutant embryos.

While cell detachment from the dorsal ectoderm that accompanies NB delamination and enables the basal positioning of the first born NBs may depend on a reduction of cell-cell adhesion, the changing positions of NBs and their derivatives along the DV axis may be influenced or maintained by cell interactions with the ECM. Consistent with this possibility, the distribution of the major Drosophila beta integrin subunit, βPS, and the ECM ligand, Laminin, visualised using GFP tagged transgenes revealed spatiotemporally regulated changes. Both proteins were enriched apically in the dorsal ectoderm and at the neuroepithelium-mesoderm boundary in embryos at mid-GBE to early GBR stages (Fig. 5 A, B). At the neuroepithelium-mesoderm boundary, both proteins exhibited a punctate/stippled pattern at mid GBE. The puncta became more intense with time, concomitant with increasing cell numbers and compaction within the neuroepithelium. While laminin puncta were largely restricted to the boundary, puncta rich in integrins were found arranged in at least two rows close to the boundary with the mesoderm, and also at other levels within the neuroepithelium (Fig. 5A). Laminin but not integrin distribution at the boundary became markedly continuous at early GBR (Fig. 5B). Also, while NBs contacted this boundary at mid GBE (Fig.5 A1, B1), GMCs occupied that position by early GBR. (Fig. 5, A3, B3). In the dpn+ NBs, intracellular enrichments of laminin were evident (Fig. 5 B3). These observations hint at the possibility that adhesion and deadhesion to the ECM through localised punctate, spot like adhesions might modulate the positions of the NBs and GMCs after delamination and govern the formation of layers. The continuous laminin deposition observed later might serve as a corset to confine and shape the nerve cord. A detailed assessment of the temporal correlations between morphometric states and the subcellular distribution patterns of integrins and laminin will require the simultaneous examination of the two in real-time. To determine whether cell-substrate adhesion influenced the early patterning of the neuroepithelium, we examined the consequences of removing the βPS integrin receptor on cellular organisation, fate specification and the stratification of the embryonic neuroepithelium.

### Defects in cellular organisation in the dorsal ectoderm in integrin mutants

Zygotic mutants in the βPS and αPS3 integrin subunits have previously been shown to cause defects in the patterning of axon tracts and in the condensation of the ventral nerve cord (Meyer et al., 2014; Stevens & Jacobs, 2002). To determine whether integrin interactions with the ECM govern the organisation of the early embryonic neuroepithelium that we described above, we generated embryos that lack both maternal and zygotic (mz) contributions of the βPS integrin subunit by obtaining embryos from germline clones of the amorphic *myospheroid^1^* allele (*mys^1mz^*). In a fraction of fixed *mys^1mz^* embryos the apical surface of the dorsal ectoderm was not flat and was thrown into folds (Fig. 6 B1, B3) compared to the heat-shocked controls (Fig. 6 A1, A3). Additionally, the cells of the dorsal ectoderm in the mutants appeared less columnar (Fig. 6 A4, B4, C2, C2’, D2, D2’) and showed cellular patches with markedly reduced levels of E-Cadherin (Fig. 6 E1-4, F1-4). They also showed a significant and widespread downregulation of discs large (dlg), a key component of the basolateral polarity complex localised to septate junctions (Fig.6 C-F). Real-time confocal microscopy of the dorsal ectoderm in mutant embryos expressing endogenously GFP-tagged E-Cadherin (shg GFP) revealed the progressive severity of E-Cadherin loss from subapical cell membranes (Fig.6 G, H). These observations uncover a role for integrins in maintaining cellular organisation and cell-cell adhesion in the dorsal ectoderm.

### Defects in stratification of the neuroepithelium in integrin mutants

The underlying neuroepithelium containing the NBs and their derivatives was also affected in integrin mutants. In contrast to the layered organisation observed in control embryos at Stage 12 (Fig. 6 A3, A4), the neuroepithelium in *mys^1mz^* embryos was disorganised. Its thickness was non-uniform, with anywhere between one and seven cell rows, as evident from the orthogonal and sagittal slices (Fig. 6 B3, B4). Cells within the neuroepithelium were not as compactly packed as in control embryos as evident from the presence of cellular aggregates and acellular gaps in some regions (Fig. 6 B3). In addition to the gaps within the neuroepithelium, gaps between it (the basally positioned pros positive cells) and the underlying mesoderm (pros negative) were also evident in a significant fraction of mutant embryos in marked contrast to the tight apposition observed between these two tissues in control embryos (∼50%, n=5; Fig. 6 A3-A4, B3-B4, 7K). The mesodermal layer also looked less continuous compared to controls (Fig.6 A2-A4, B2-B4).

Both neuroepithelial stratification and detachment defects coexisted in approximately 60% (n=50) of *mys^1mz^* embryos, and were also observed in a fraction of embryos lacking only maternal βPS (*mys^1m^*) (Fig. 7K). These results highlight a requirement for both maternal and zygotic integrin in maintaining the organisation of the dorsal ectoderm and in patterning the layered organisation of the neuroepithelium. Whether the defects in stratification are a result of defects in cellular organisation and cell adhesion, or whether specific integrin-ECM interactions between cells within the neuroepithelial layer guide its stratification remains to be resolved.

### Mispositioning of NBs and GMCs along the dorsoventral axis of the neuroepithelium in integrin mutants

The defective neuroepithelial stratification in *mys^1mz^* embryos described above was most evident in the altered distribution of NBs and GMCs which were no longer arranged in well-defined dorsoventral rows. In *mys^1mz^* mutant embryos, dpn+ NBs and pros+ GMCs were found intermixed across the neuroepithelial layers (Fig. 7 B2, D2). Remarkably, dpn^+^ NBs were also found in the dorsal ectoderm (surface) in more than half of the *mys^1mz^* embryos (n=50; Fig. 6D, 7K), in stark contrast with control embryos, in which dpn^+^ NBs are found in progressively more basal locations and rarely in the ectodermal layer (Fig. 6C, 7 A1, A2, B1, B2). Approximately half of the total dpn+ cell population in *mys^1mz^*embryos (including those on the surface and in the immediately basal layer) were found in the dorsal ectoderm (Fig. 7I). Deadpan positive cells were also found in the dorsal ectoderm in embryos in which cellular disorganisation was not so pronounced (Fig. 6D2’). Prospero positive GMCs, like the dpn+ NBs, were also variably distributed throughout the neuroepithelium, including both the surface and basal layers (Fig. 7J) with approximately 50% (n=50) of mutant embryos exhibiting surface-localised pros+ cells. Prospero positive cells accounted for ∼10% of all cells in the dorsal ectoderm and ∼50% of cells in the basal layer immediately underlying the ectoderm. No pros+ cells were detected in either of these two layers in control embryos where they were found in deeper layers (Fig. 7J). Hunchback (hb) positive cells mark the first born NBs and occupy progressively more basal locations in the neuroepithelium. Like dpn+ and pros+ cells, hb+ NBs were also found in the dorsal ectoderm in *mys^1mz^* embryos during early stages and haphazardly distributed within the neuroepithelium at later stages (Fig. 7 E-H). While we have not determined whether the absolute numbers of NBs or GMCs is altered in integrin mutants, our findings suggest that the expression of the fate determinants does not depend on integrins, and that delamination is not necessary for NB differentiation. Instead, our observations uncover a role for integrins in influencing the position and possibly also the timing of differentiation of NBs and their derivatives.

### Altered morphology and spindle orientation in integrin mutant NBs

We surmised that the presence of NBs and GMCs in the dorsal ectoderm might reflect differentiation and division without delamination, or their reintegration into the ectoderm after delamination. The dpn+ cells in the dorsal ectoderm in integrin mutants exhibited altered morphologies compared to both control ectodermal cells and to dpn+ cells in control embryos. Integrin mutant dpn+ cells were rounded compared to their dpn-neighbours in contrast to dpn+ cells in control embryos which exhibited features of one of three morphometric states S4-S6. Deadpan positive NBs in *mys^1mz^* embryos were round and lacked the distinct apical and basal features of the S4 morphometric state, characterised by marked apical constriction, basal expansion and detachment of its apical end from the ectoderm. They were instead integrated into the ectodermal layer (Fig. 6 C, D, 7 A2, B2, E4, F4, G4, H4).

NBs in control embryos divide after delamination orthogonally to the plane of the epithelium with the spindle oriented along the apicobasal axis, and asymmetrically to generate a larger NB (dpn+) and a smaller GMC (nuclear pros+). This asymmetry is marked prior to division by a basal cortical crescent of Prospero. The division plane is easily identified by spindle position and DNA organization. In addition, as described earlier, the subcellular distribution of dpn protein provides a readout of its cell cycle stage, being diffuse and cytoplasmic in metaphase, and compact and nuclear in telophase (Fig. 8A, 4 D2).

**Figure 8:**
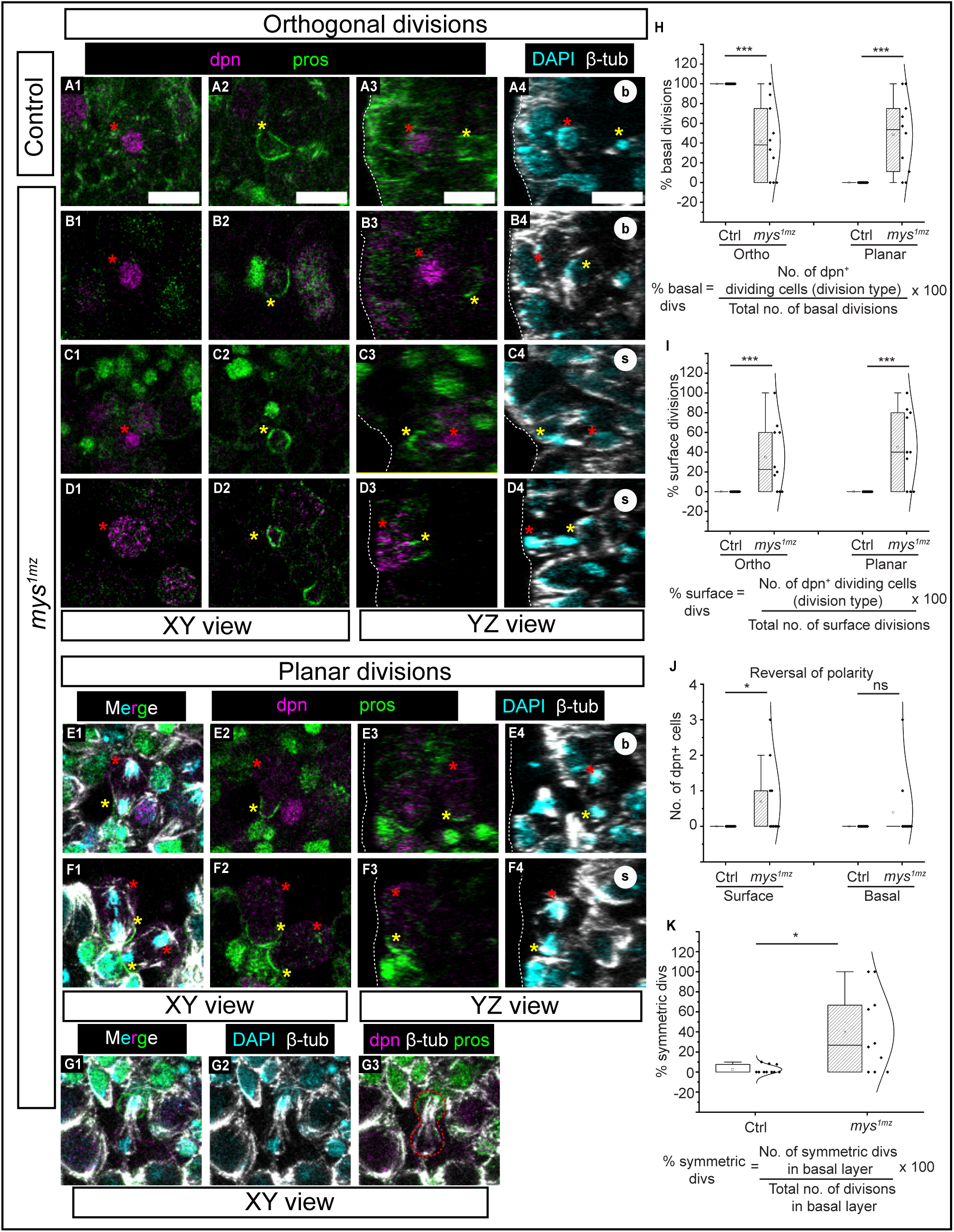
Defects in spindle orientation in dividing neuroblasts in integrin mutants. Change fig Representative images showing XY and YZ views (as labelled) from control (A) and *mys^1mz^* mutant (B-G) embryos immunostained for β-tubulin (white), deadpan (magenta), prospero (green) and DAPI (cyan). Neuroblasts dividing either orthogonal to the epithelial plane (A-D) or within the epithelial plane (E-G) are shown. The divisions occur either basally (A, B, E) or in the surface (C, D, F). NB division with an apparent reversal of polarity (C) or occurring symmetrically (G, red dotted line) are shown. White dotted lines indicate the apical surface of the neuroectoderm. Red and yellow asterisks indicate neuroblast and GMC respectively. Scale: 10 μm. Percentage/ relative frequency of basal (H) or surface (I) divisions occurring either orthogonally or in plane in control and *mys^1mz^* mutants. (J) Quantification of dpn+ NB showing reversal of polarity at the surface and basally in control and *mys^1mz^*mutants. (K) Percentage/relative frequency of symmetric divisions in the basal layer in control and *mys^1mz^* mutants. In the box plots in H-K boxes show median (horizontal line) ± interquartile range; the mean is indicated by ‘□’ and the sample size is in brackets. (H, I) Student’s t-test or (J, K) Mann-Whitney test was used for statistical analysis (*-p < 0.05, **-p < 0.01, ***-p < 0.001, and ns-not significant).

**Figure 8:**
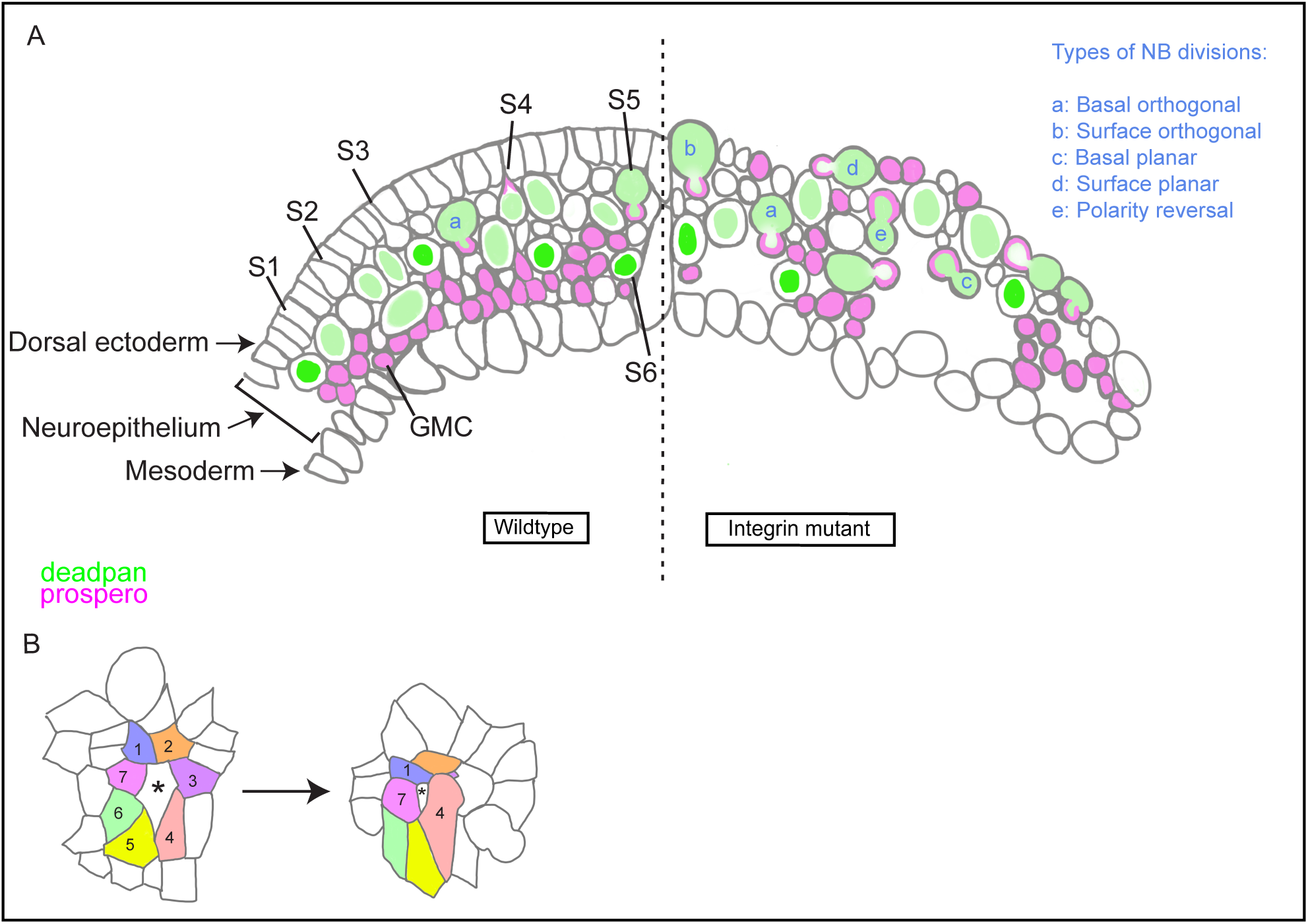
Consequences of the loss of integrins on neuroepithelial organisation, cell morphodynamics and division (A) Schematic representation of the organisation of the dorsal ectoderm, the underlying neuroepithelium and the mesoderm to which it is closely apposed in control embryos (left half) showing the morphometric states (S1-S6) accompanying neuroblast specification and division and their associations with the expression of fate determinants Dpn (NB, green) and Prospero (GMC, magenta). The types of division are indicated by the small case alphabets. In integrin mutants (right half), neuroepithelial disorganisation, progenitor type intermixing and mispositioning, as well as altered positions of the division plane are observed. (B) An apical view of a cell cohort containing a NB (asterisk) showing a radially asymmetric rosette. The dynamic neighbourhood of the NB is characterised by nearest neighbour losses. Nearest neighbours are marked by filled colours.

Deviations in division patterns were observed in *mys^1mz^*mutants. Deadpan positive cells in the dorsal ectoderm (surface) in the mutant embryos were found to divide orthogonally and asymmetrically as in control cells (∼30% of surface divisions, n=10; Fig. 8 A, B, J, K) as well as asymmetrically in the plane of the ectoderm (∼40% of all surface divisions, n=10; Fig. 8, C, F, K). This was also true of dpn+ cells located in the basal layer in which 41% of divisions occurred orthogonally and ∼50% in the plane of the epithelium n=10; Fig. 8I). Some dpn+ NBs (Fig. 8J) exhibited a polarity reversal with a dorsal Prospero crescent, and others (40%) exhibited division that yielded dpn+ and pros+ cells that were symmetric in size (Fig. 7 A1, A2, B1, B2, K). These observations reveal that while the division of the NB and its asymmetry is unaffected by the loss of integrins, the division plane is sensitive to integrin loss. Whether the latter is due to the defects in cell adhesion that remove rotational constraints or whether integrins alter spindle positioning remains to be resolved. The defects, we believe, could nonetheless contribute to the mispositioning of NBs and their derivatives within the neuroepithelium. Real-time confocal microscopy with fate determinant gene and morphodynamic markers should enable the delineation of the altered morphodynamics of NB delamination and division and the acquisition of fate in integrin mutants. What the consequences of NB repositioning in early neurogenesis are on patterning the stereotypical circuitry of the embryonic and larval nervous system remains to be resolved.

## Discussion

In the work we describe above, we used NB delamination as a test case to examine the spatial, temporal and molecular hierarchies in the link between the morphodynamics of delamination, the acquisition of cell fate and the formation of a stratified neuroepithelium. Our investigations have uncovered temporal correlations between the morphometric state of a delaminating neuroblast and the time of appearance and subcellular distribution of fate determinant proteins. Our observations suggest a transcription independent and morphology dependent mechanism by which progenitor fate and position can be regulated in a morphogenetically active epithelium. The differences we uncover between NB delamination and delamination elsewhere lend support to the possibility that the morphogenetic activity of an epithelium can influence the nature of cellular morphodynamics. Our results also underscore the importance of the regulation of NB microenvironment provided by dynamic integrin-ECM interactions and suggest their role as placeholders and possibly also timekeepers in patterning the embryonic neuroepithelium. Together, our work provides a glimpse into the emergence of the stereotypically layered organization of the nervous system from the delamination of single cells (Fig. 9).

### Distinct morphodynamic features of delaminating NBs: long stalks with bulbous bases

Our analysis of NB delamination in 4D has uncovered several morphological features that go beyond the anisotropy in apical constriction and junction loss previously reported (Simões et al., 2017). Two features, progressive constriction along the lateral membrane and the bulbous expansion of the basal region make the shape of the delaminating NB markedly different from the frustum/prism shapes observed in apical constriction (Leptin, 1999; Martin et al., 2009). The long stalk and the long duration to complete delamination might facilitate the timely and accurate basal positioning of the neuroblast and its differentiation (the expression of the NB fate determinant, dpn) by ensuring long lasting apical adhesive contacts with its neighbours. The bulbous basal expansion likely reflects basal relaxation or mitotic rounding prior to the detachment of the NB from the ectoderm and enables asymmetric cell division shortly thereafter. These speculations are consistent with our observations that NB differentiation and position are sensitive to adhesion (see below) and that cell division ensues within minutes of detachment of the NB from the surface ectoderm. Subcellular perturbations will be necessary to elucidate the roles of apical and basal contacts in the positioning of progenitors. Our results reveal that the morphodynamics of NB delamination bears some resemblance to the morphodynamics of delaminating mouse neural crest cells (Moore et al., 2024). While neural crest cells migrate after their delamination, NBs reposition themselves along the thickness of the neuroepithelium using subcellular mechanisms that bear some similarities to progenitor positioning in the zebrafish Kupfer’s vesicle (Pulgar et al., 2021). The morphodynamic transitions we observe during the delamination of embryonic neuroblasts differs from the collective delamination of the Drosophila mesoderm during invagination of the ventral furrow. Mesoderm cells exhibit a sequence of shape transitions that include (pulsed) apical constriction and cell shortening to create frustum shaped cells that maintain contacts (albeit weakened) with their neighbours and then flatten or divide after invagination is complete (Leptin, 1999; Martin et al., 2009; Seher & Leptin, 2000). What factors guide the differential cortical mechanics along the apicobasal axis of ingressing/delaminating cells and what functions they serve remain to be resolved.

### Delamination in a dynamic neighbourhood: implications for lateral inhibition

Our observations uncovered marked changes in the number of nearest neighbours around a delaminating neuroblast, and reduced collectivity/supracellularity in their behaviour, leading to the lack of radial symmetry in the NB rosette. This is contrast with delamination in the amnioserosa in which the predominantly isotropic constriction of the delaminating cell, and collective anisotropic shape changes in the nearest neighbours (characterized by the shortened interfaces with the delaminating cell and the lengthened interfaces between the nearest neighbours; Meghana et al., 2011) resulted in radially symmetric rosettes. We speculate that these differences must have their origins in the prevailing tone of morphogenetic activity in the epithelium. Amnioserosa cells during dorsal closure are programmed to undergo apical constriction, that possibly accounts for both the higher rate of constriction and the formation of more symmetric rosettes (Meghana et al., 2011; Saravanan et al., 2013; Solon et al., 2009; Toyama et al., 2008). In contrast, NB delamination occurs in the backdrop of germband extension and retraction, with cells programmed to remodel junctions or undergo division (Bertet et al., 2004). The dynamic heterogeneity in the NB neighbourhood could contribute to rosette asymmetry. We also uncovered heterogeneities in the rate of constriction in delaminating NBs. While some NBs constricted rapidly, and with rates similar to delaminating amnioserosa cells (3.6 μm²/min; Meghana et al., 2011) many NBs did so at rates that are on average between two and three-fold lower. These differences may also be influenced by the neighbourhood which has been shown to provide forces for and govern the dynamics of delamination in amnioserosa (Meghana et al., 2011; Saravanan et al., 2013). The dynamic heterogeneity in the NB neighbourhood raises the question of how NB numbers and position are maintained, and how signals including Notch whose interactions with the ligand Delta promote lateral inhibition that NB specification depends on are integrated. Our observations that nearest neighbours can delaminate in the neuroectoderm suggest that the mechanisms and outcome of lateral inhibition in a dynamic epithelium may differ from that in a static epithelium. Notch signalling has been shown to depend on cell and interface geometry (Shaya et al., 2017) and on morphogenetic activity (Falo-Sanjuan & Bray, 2022) and it is tempting to speculate that dynamic changes in signal delivery or reception could be both responsive to and influence cell morphodynamics.

### The link between morphodynamics and fate acquisition: subcellular localisation of fate determinants

Fate in developmental biology is traditionally scored on the basis of expression of the protein products of ‘fate determinant’ transcription factors or their target genes. Recent methods have relied on the delineation of cell type specific transcriptomes (Mircea & Semrau, 2021). Our observations with antibodies against protein products of bona-fide NB (Deadpan) and GMC (Prospero) fate determinants uncovered heterogeneities in their subcellular distribution that bore correlations with distinct morphometric states. A common feature of both proteins was their existence outside the nucleus. Dpn was first detected in the nucleus in a fraction of NBs in the S4 state, characterised by complete apical constriction and detachment of the NB from the dorsal ectoderm. The subsequent diffuse and cytoplasmic distribution of dpn in S5 likely indicates nuclear envelope breakdown in preparation for mitosis. The GMC determinant Prospero was enriched in the apical cortex of the NB prior to its detachment and to its enrichment at the basal cortex before it was detected in the nucleus of the GMC. These correlations suggest that the crosstalk between morphodynamics and fate specification may operate through the post-translational regulation of protein distribution. Indeed, many fate specification principles including Notch and β-catenin show dynamic subcellular distribution patterns that can also be modulated by mechanical forces (Farge, 2003; Meloty-Kapella et al., 2012; Mitrossilis et al., 2017).

### Integrins as integrators of morphodynamics and fate specification: placeholders and timekeepers

Our observations identified dynamic changes in the distribution of integrin and laminin during the delamination of the neuroblast. Both localised to and became progressively enriched in puncta that decorated the neuroepithelium-mesoderm boundary from the beginning of neurogenesis (mid GBE), and the distribution of laminin at the boundary became continuous by mid GBR. Remarkably, the distance between the apical surface of the dorsal ectoderm and the mesoderm boundary remained relatively constant. These findings suggest that the progressive dorsoventral expansion of the neurectoderm occurs in a confined space and is accompanied by changes in cell packing densities. We speculate that this necessitates that the neurectoderm-mesoderm boundary be ‘stiff’ enough to prevent expansion but dynamic to allow compaction and the formation of layers. Consistent with this possibility, the complete absence of the βPS integrin resulted in detachment between the mesoderm and the neuroepithelium, marked heterogeneity in its thickness, and cellular disorganisation within it.

Puncta containing βPS integrin marked two rows, including the neuroepithelium-mesoderm boundary and were also found at different positions within the neuroepithelium suggesting local, possibly dynamic contacts between cells in different layers through ‘spot’ adhesions, that matured to more ‘stable’ adhesions as layers formed. The presence of intracellular laminin in NBs also suggests that integrin ECM interactions may be dynamically regulated through the regulation of ECM secretion. Consistent with this possibility, NBs and GMCs were mispositioned and intermixed across the neuroepithelium in the absence of integrins. Their specification rules out the requirement of integrins for fate determinant gene expression. What was striking however was their altered shapes and positions, notably their presence in the dorsal ectoderm. Whether this is indicative of a failure in or altered direction of delamination, or an increased propensity for reintegration, or whether the constitutive loss of adhesion to the substrate triggers precocious differentiation into a NB remain to be determined.

The loss of integrins also affected lateral adhesion, evident from the reduction in E-Cadherin and discs large, and the plane of asymmetric cell division, without affecting division asymmetry. These observations suggest that the loosening of lateral cell contacts in the absence of integrins might lift the constraints that time NB differentiation and maintain neuroepithelial organisation. Integrins have been shown to influence cell-cell adhesion and to alter delamination dynamics (Meghana et al., 2011; Narasimha & Brown, 2004). Lateral adhesion through adhesion molecules at the septate junction have been shown to enable the restoration of epithelial organisation by promoting reintegration of cells misplaced due to altered division planes (Bergstralh et al., 2015). Integrins have also been shown to be necessary for the maintenance of monolayers, and its absence causes ‘multilayering’ in the Drosophila ovarian follicle epithelium (Fernández-Miñán et al., 2007). Our observations raise the possibility that integrin adhesion, through focal contacts or through the regulation of lateral adhesion maintains neuroepithelial progenitor position and stratification. The molecular mechanisms underlying their requirement remain to be discovered.

Our work also uncovers an early requirement for integrins in the patterning the neuroepithelium: positioning the progenitors, and possibly also timing their differentiation, thus functioning as place holders and time keepers. Drosophila integrins have previously been shown to pattern axon tract fasciculation and ventral nerve cord condensation later in embryogenesis (Stevens & Jacobs, 2002) and whether these defects might also have their origins in the early phenotypes we observe remains unknown. The maintenance of stem cell niches in the mouse neocortex also depends on integrin function, likely through its influence on intrakinetic nuclear migration that contributes to the layering of other neuroepithelia (Loulier et al., 2009). What the nature of the dynamic interactions of integrins with the ECM are, and what subcellular and molecular mechanisms underlie the influence of integrins on progenitor specification and positioning in the Drosophila neuroepithelium remain to be discovered. Our observations bear some similarities to a recent study that demonstrated the requirement for reduced-integrin ECM interactions to promote germline entry/germ cell differentiation in the mouse gonad (Makhlouf et al., 2024).

### Building cell layers: morphodynamic and evolutionary diversity

Across development and evolution, multiple morphodynamic strategies enable the formation of layers. NB delamination in Drosophila, mesoderm internalization in Chironomous, border cell specification in the Drosophila ovary and vertebrate neural crest cell delamination all involve the delamination of single cells (Gouignard et al., 2018; Montell, 2006; Urbansky et al., 2016). In contrast, the formation of the Drosophila mesoderm and the vertebrate neural tube rely on the collective ingression or delamination of a cell sheet (Leptin, 1999; Schäfer et al., 2014; Yamaguchi & Miura, 2013). What morphodynamic, cytoskeletal and adhesion changes distinguish the two types of delamination, and how these differences contribute to the efficiency or robustness of tissue patterning will be interesting to discover.

### Conclusions

Our work throws light on the asymmetric organization of rosettes during the specification of neuroblasts and challenges the idea of how lateral inhibition might work in a constantly changing neighbourhood. It provides insights into how organized layers can be built from single cells and suggests that integrin-ECM interactions serve as placeholders and timekeepers during the patterning of the stratified Drosophila embryonic neuroepithelium. Our work also uncovers an earlier requirement of integrins during embryonic neurogenesis. Understanding the subcellular dynamics of integrin-ECM interactions and the mechanisms by which they influence and integrate cell morphodynamics, fate and position will be interesting avenues to pursue.

### Limitations of the study

While our work has established temporal correlations between morphodynamic states and fate determinant protein distribution in fixed embryos, a real-time analysis of these correlations including an examination of the temporal patterns of transcription will enable a better definition of the temporal hierarchies.

Our work has also not established a causal relationship between cell shape change and fate determinant protein distribution. This will necessitate the examination of fate determinant gene expression in genetic perturbations targeting regulators of specific aspects of cell shape changes.

Our work identifies adhesion to ECM as a regulator of both cell morphodynamics and neuroepithelial patterning. While the distribution patterns of integrin and ECM, and the phenotypes observed in germline clones provide clues about the nature of their requirement, a real-time analysis of the dynamics of integrin and ECM as well as tissue specific perturbations and rescue of integrin function will be necessary to establish their precise or direct roles in neuroepithelial patterning.

Our study does not resolve the mechanistic basis for the requirement of integrins.

## Supporting information

Supplementary File with text and figures

## Acknowledgements

We thank Nick Brown, Tadashi Uemura, the Bloomington Drosophila Stock Centre (BDSC), the Vienna Drosophila Resource Centre (VDRC) and the Developmental Studies Hybridoma Bank (DSHB) for stocks and reagents, and members of the MN lab for discussion. We acknowledge support from TIFR/DAE, India (RTI4003 to MN and graduate fellowships to LD and AG) and from the DG Bhosale Fellowship (to LD). MN wishes to dedicate this work to the memory of Prof. Veronica Rodrigues.

## Author contributions

Lamiya Dohadwala: Formal analysis; Investigation; Visualization; Methodology; Writing – original draft; Writing – review and editing; Performed all experiments, data generation and statistical analysis.

Anupriya Garg: Formal analysis; Visualization; Writing – review and editing; Contributed data generation and analysis

Maithreyi Narasimha: Conceptualization; Resources; Formal analysis; Supervision; Funding acquisition; Visualization; Methodology; Writing – original draft; Writing – review and editing; Conceived project, designed experiments and methodology, wrote first draft with LD, revised and edited manuscript, supervised the project and acquired funding.

## Materials and Methods

### Materials

#### Fly stocks

All fly stocks used are listed in the table below.

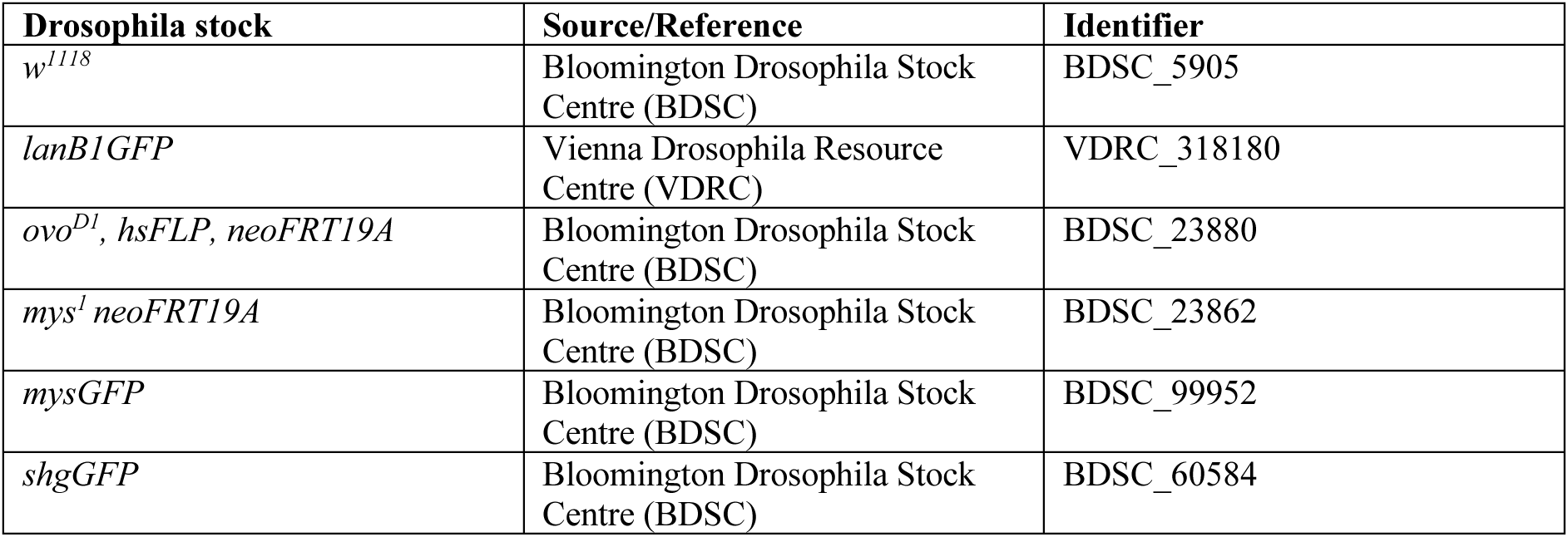

#### Antibodies and reagents

| Primary antibody | Animal raised in | Dilution | Identifier, Source |
| --- | --- | --- | --- |
| E-Cadherin | Rat | 1:10 | DCAD2, DSHB |
| GFP | mouse | 1:2500 | sc-9996, Santa Cruz |
| GFP | Rabbit | 1:2500 | A11122, Invitrogen |
| discs-large | Mouse | 1:20 | 4F3, DSHB |
| deadpan | Rat | 1:10 | ab195173, Abcam |
| prospero | Mouse | 1:10 | 1D4, DSHB |
| β-tubulin | Rabbit | 1:1000 | ab179513, Abcam |
| Hunchback | Mouse | 1:1000 | ab197787, Abcam |
| βPS integrin | Mouse | 1:10 | CF.6G11, DSHB |

DSHB: Developmental Studies Hybridoma Bank

| Secondary antibody | Dilution | Identifier, Source |
| --- | --- | --- |
| Goat anti-mouse 488 | 1:200 | Invitrogen, A11029 |
| Goat anti-rabbit 488 | 1:200 | Invitrogen, A11034 |
| Goat anti-mouse 568 | 1:200 | Invitrogen, A11031 |
| Goat anti-rabbit 568 | 1:200 | Invitrogen, A11036 |
| Goat anti-rat 647 | 1:200 | Invitrogen, A21247 |

| Stain | Dilution | Identifier, Source |
| --- | --- | --- |
| Rhodamine Phalloidin | 1:100 | Invitrogen, R415 |
| DAPI | 1:1000 | Sigma D9542 |

#### Software/ Codes

| Software and Plugins | Source or reference | Related weblink |
| --- | --- | --- |
| ImageJ | National Institute of Health (Schindelin et al., 2012) | https://imagej.nih.gov/ij/ |
| Tissue Analyzer | (Aigouy et al., 2016) | https://github.com/baigouy/tissue_analyzer |
| OriginPro 2021 | Origin Software | https://www.originlab.com/origin |

## Methods

### Embryo collection

Flies were allowed to lay eggs on standard cornmeal agar fly food for 5-6 hours at 29°C to enrich for germband extension stages. For live imaging, embryos were dechorionated for 2 minutes in 50% bleach diluted in water. They were then collected using a Corning Cell Strainer (100μm pore size) and rinsed under tap water to remove residual bleach. Embryos were handled using a paintbrush and mounted on 22mm X 40mm coverslips (0.15-0.17mm thickness) which were coated with a layer of Halocarbon 27 oil (Sigma Life Sciences). Imaging was performed using an inverted confocal microscope (Olympus FV3000).

### Immunofluorescence

For immunostaining, embryos were harvested and stained using standard protocols (Narasimha & Brown, 2006). The specific protocol used in this study is as follows: after dechorionation (as described above), embryos were fixed in a freshly prepared fixative solution containing a 1:1 ratio of heptane and 4% paraformaldehyde in 1X PBS. Fixation was carried out in a 2mL micro-centrifuge tube (MCT) placed on a rotor (Rotospin, Tarsons 3071) for 20 mins at 23rpm at room temperature (RT). Fixed embryos collected at the interphase between the aqueous and organic phases, the solution was carefully removed and replaced with 1:1 ratio of heptane and methanol. The solution was vortexed for 2 mins to remove the vitelline membrane. Devitellinized embryos settled at the bottom of the tube, after which the solution was carefully removed, and the embryos were washed thrice with methanol and stored in methanol at 4°C for future immunostaining. For phalloidin staining, embryos were devitellinized in 90% ethanol instead of methanol, followed by three washes with absolute ethanol and stored at 4°C.

For immunostaining, methanol or ethanol was removed, and embryos were subjected to four gravity washes in PBT (0.3% TritonX-100 [v/v] in 1x PBS), followed by a 30 mins wash on the rotor at RT. Embryos were then blocked in PBTB (0.5% BSA [w/v] in 0.3% PBT [v/v]). After blocking, the solution was removed, and embryos were carefully transferred into 0.6mL MCTs, to which the primary antibody cocktail (prepared in PBTB) was added. Embryos were incubated for 16hrs on the rotor at 4°C. Following primary incubation, the antibody solution was removed, and embryos were subjected to five quick gravity washes with PBT, followed by three 20mins washes on the rotor at RT. The secondary antibody cocktail (prepared in PBTB) was then added and incubated for 2hrs on the rotor at RT. After secondary incubation, embryos were given 5 gravity washes, followed by four washes on the rotor for 40, 20, 10 and 10mins, respectively. For DAPI staining, embryos were incubated with DAPI (1:1000 in PBT) during the 40 mins wash. After completing all washes, embryos were stored in Vectashield mounting medium (Vector Laboratories, H-1200-10) at 4°C and imaged the next day once embryos had settled in the medium.

### Generation of *mys^1^* germline clones

Maternal and zygotic mutants of the βPS integrin mutant allele *mys^1^* (Bunch et al., 1992) were generated by making germline clones using the FLP/FRT system (Chou & Perrimon, 1992). Females of genotype *mys^1^neoFRT19A/FM7cKrGFP* females were crossed to *ovo^D1^, hsFLP, neoFRT19A/Y* (BDSC_23880) males and allowed to lay eggs for 24hrs at 29°C on standard cornmeal agar fly in glass vials. Embryos were aged for 48hrs to the second instar larval stage (when primordial germ cells divide), then heat shocked in a water bath at 37°C for 1.5 hrs to induce FLP-mediated site-specific recombination under the hsp70 promoter. Vials were returned to 29°C until eclosion.

Female progeny of genotype *mys^1^ neoFRT19A/ovo^D1^, hsFLP, neoFRT19A* were identified by the absence of the Bar-eyed marker on the FM7c balancer. Successful mitotic recombination was confirmed by the presence of wing blisters, a phenotype characteristic of integrin loss (Prout et al., 1997). These females produced germ cells of three genotypes: *mys^1^/mys^1^*, *mys^1^*/*ovo^D1^*and *ovo^D1^/ ovo^D1^*. As *ovo^D1^* is a dominant female-sterile mutation, only *mys^1^/mys^1^* germ cells could give rise to fertilized eggs, enabling selection of maternally null mutants.

Heat-shocked females were crossed to FM7cKrGFP males. All resulting embryos were maternally null for *mys^1^*, with FM7cKrGFP used to distinguish maternal-zygotic (*mys^1^/Y*) from maternal-only (*mys^1^/FM7cKrGFP*) mutants.

### Genotypes examined: see Supplementary File

Contains a figure-wise list of genotypes examined.

### Image acquisition

All images were acquired on an Olympus FluoView3000 laser scanning confocal microscope at room temperature. Low-magnification images were taken using 10X/0.4 NA or 20X/1.4NA oil objectives. High-magnification 3D images of immunostained embryos were acquired using a 60X/1.42 NA oil objective (at 1.0-2.5X digital zoom) and a Z-step size of 0.41μm at scanning speeds of 4 or 8μs/pixel. For live imaging of embryos carrying the shgGFP transgene, optical sections were acquired at a Z step size of 0.8μm to a depth of 40μm into the embryo at a scan speed of 2μs/pixel.

### Post-acquisition image processing

No manipulations other than level adjustments to stretch the intensity histogram (when necessary) were applied. Adobe Photoshop and Adobe Illustrator were used for compiling of the images and graphs.

### Defining the ROI for analysis

Using the dorsal midline as the positional reference, we imaged three to four rows of cells starting from and running parallel to the dorsal midline and quantitatively tracked the morphodynamics of delaminating NBs, which progressively abutted the dorsal midline to become the medial rows of NBs under the epithelial sheet. The anterior-posterior boundary of the ROI considered for morphodynamic analysis roughly spanned columns 3-7 of achaete expression, which marks the proneural clusters arranged in stereotypic rows and columns within the germband during stages 8-9 of embryogenesis (Fig. 1C1). A rectangular ROI of dimensions 63.75 x 49.7 μm was centred at the dorsal midline on the germband, with the left bound of the ROI at its anterior most tip was used in Fig. 1, C-E and Fig. 4. For all quantifications from fixed preparations, in Figs. 6-8, a rectangular ROI of dimensions 120 x 90μm centred at the dorsal midline was used.

### Defining the temporal window for analysis of cellular morphodynamics

NBs delaminate from the ectoderm in five successive waves. NBs that delaminate during the second and third waves, which occur at stages 9 and 10 of embryogenesis, when the germband length is 58-65% of the egg length, were chosen for analysis. The extent of germband elongation (GBE) in five embryos was measured from low magnification time-lapse movies of embryos carrying the SquashUtrophinGFP transgene. The fully extended germband was designated as time 0 (t0), and measuring its length relative to the egg length retrospectively enabled us to extrapolate the time required to reach the fully extended stage from any initial stage. Embryos with relative germband lengths (55-70% EL) that captured the second and third waves and ensured at least a 60-minute time window for tracking a delaminating NB was used. For fixed preparations, embryos having germband length 68-70% of E.L, capturing 2^nd^ and 3^rd^ waves of neurogenesis were chosen for analysis (Fig. 1 A, B, F, 2 A-H, 3 A-H, 6G, H).

### Cellular morphodynamics analyses

For the analysis of apical cell area dynamics, maximum intensity projections of optical slices from 2.4 μm to 3.2 μm depth from the vitelline membrane, for each time frame were generated. Tissue Analyzer was used to automatically segment cell outlines in time-lapse movies from these stacks. To measure the apical area and aspect ratio of a single NB or its NN, the NB or NN was manually selected at the earliest time point that the cell of interest could be tracked. Manual corrections of cell outlines were applied when necessary. Apical areas and aspect ratios of 28 neuroblasts and its cohorts from five embryos were temporally aligned by defining the time point at which their apical area first fell below 3 μm^2^ as t0. Cell area dynamics were analyzed by plotting the changes in apical area as a function of time (Microsoft Excel).

To determine the morphodynamic state of delaminating NBs, cells were optically sliced along the YZ axis to visualize the 3D morphology of the cells. The cell shapes were classified into six morphometric states by measuring the ratios of the height and width of the stalk and bulb, determining their position within the epithelium (either within the epithelial sheet or basally), and observing the formation of a smaller basal bud, which indicates the asymmetric division of the neuroblast into a neuroblast and a ganglion mother cell (GMC).

NBs were classified as undergoing expansion prior to net constriction if the duration of increase in area prior to that start of constriction was at least 5 minutes, expansion occurred within 60 minutes prior to the completion of delamination, and had a fold change of area that was greater than or equal to 1.5 (Fig. 2 A, B, D, E).

### Cell counting and division orientation

The number of nuclei (marked by DAPI) were used to count the total number of cells in the neuroepithelial layers. Dpn, Pros and tubulin/DAPI were used to identify NBs, GMCs and spindle orientation patterns respectively. In this study, the basal layer refers to the cells immediately beneath the surface ectoderm. Only the surface and basal layers were quantified for the analysis of relative frequencies of cell division planes (Fig. 8), or the relative distribution of NBs and GMCs in each layer (Fig. 7). In metaphase and anaphase neuroblasts, orthogonal divisions are defined as spindle orientations perpendicular to the plane of the epithelium (along the apicobasal polarity axis), and planar divisions are defined as spindles oriented within the plane of the epithelium (perpendicular to the apicobasal axis).

### Graphs and statistical analysis

All graphs were prepared, and all statistical analyses were performed using OriginPro 2021. Box and whisker plots, bar graphs and line graphs have been used for data visualization in this study. Box plots (used for visualising the distribution of neighbour dynamics and cell type heterogeneity in Figs. 3 D-F, 3H,4C, 7I, 7J, and 8H-K) display the median ± interquartile range (IQR). The thick black lines represent the median, the ‘□’ sign indicates the mean, and the box limits correspond to the 25th and 75th percentiles. Whiskers extend to 1.5 times the IQR from the 25th and 75th percentiles. Dots represent individual data points. The sample size (n) analysed for each genotype is mentioned in the graph. P-values, statistical tests used, sample sizes, and data distribution types are detailed in the Statistical Table (Supplementary File). No statistical test was used to predetermine the sample size, and no data was excluded from the analysis. The statistical significance between central tendencies of normally distributed datasets was determined using the Unpaired student’s *t*-test, while the Mann-Whitney *U*-test was used for non-normal distributed data. P values, sample sizes, normality test and the statistical test employ are provided in the Statistical table (See Supplementary File) for the datasets compared. The statistical significance is denoted as follows, *-p <0.05, **-p < 0.01, ***-p <0.0001 and ns-not significant.

### Statistical table (See Supplementary File)

Contains a figure wise list of statistical comparisons

## Figure legends

**Supplementary Figure 1.** Apical area dynamics of delaminating neuroblasts from 5 embryos. A1-A15 show three constriction rates and A16-A28, a single constriction rate.

**Supplementary Figure 2.** (A1-A28) Quantitative morphodynamic analysis of 28 delaminating NBs from 5 embryos showing aspect ratio (red) and area (black) dynamics. t0 marks the time at which the area of the delaminating cell is less than 3μm^2^. A1-A6, A7-A12 and A13-A28 show respectively a decrease, no change, or increase in shape anisotropy/aspect ratio in the 10 minutes prior to the end of delamination.

**Supplementary Figure 3.** (A1-A6) Apical area dynamics of 6 delaminating NB cohorts. Black lines indicate the delaminating NB and the coloured lines are its nearest neighbours. Some nearest neighbours show an increase in area closer to the end of delamination. Asterisks mark neighbours lost.

