## Supplementary File with text and figures for "Integrins pattern the Drosophila embryonic neuroepithelium by influencing progenitor morphodynamics, division and position"

**Supplementary resources**

**Genotypes examined**

Figure 1

A, B, F: *w*; sqh::UtrophinGFP/ sqh::UtrophinGFP

C-E: *w*$^{1118}$

Figure 2

A-H: *w*; sqh::UtrophinGFP/ sqh::UtrophinGFP

Figure 3

A-H: *w*; sqh::UtrophinGFP/ sqh::UtrophinGFP

Figure 4

A-D: *w*$^{1118}$

Figure 5

A1-A3': *y*$^{1}$, *w**, mysGFP

B1-B3': *y*$^{1}$, *w**; +/+; lanB1GFP/lanB1GFP

Figure 6

A1-A4, C1, C1', C2, C2', E1-E4: *w*$^{1118}$/Y (heat-shocked)

B1-B4, D1, D1', D2, D2', F1-F4: *mys*$^{1}$neoFRT19A/Y (maternal-zygotic mys$^{1}$ mutants)

G1-G3: *w*$^{1118}$/Y; shgGFP/+ (heat-shocked)

H1-H3: *mys*$^{1}$neoFRT19A/Y; shgGFP/+ (maternal-zygotic mys$^{1}$ mutants)

Figure 7

A1, A2, C1, C2, E1-E4, G1-G4 and Ctrl in I, J, K: *w*$^{1118}$/Y (heat-shocked)

B1, B2, D1, D2, F1-F4, H1-H4, and *mys*$^{1mz}$ in I, J, K: *mys*$^{1}$neoFRT19A/Y (maternal-zygotic *mys*$^{1}$ mutants)

*mys*$^{1m}$ in K: *mys*$^{1}$neoFRT19A/FM7cKrGFP

Figure 8

A1-A4 and Ctrl in H, I, J, K: *w*$^{1118}$/Y (heat-shocked)

B-G and *mys*$^{1mz}$ in H, I, J, K: *mys*$^{1}$neoFRT19A/Y (maternal-zygotic *mys*$^{1}$ mutants)

**Statistical table**

| Figure no. | Parameter | Populations compared | Sample size | Normal distribution | Statistical test | P-value |
|---|---|---|---|---|---|---|
| 3D | Neighbour cell area (µm²) | Start of constriction<br>End of delamination | n=83, N=5<br>n=83, N=5 | No<br>No | Mann-Whitney test | 0.00263 (*) |
| 3E | Number of neighbours | Start of constriction | n=28, N=5 | No | Mann-Whitney test | 2.8656E-10 (***) |

| | | | | | | |
|---|---|---|---|---|---|---|
| | | End of<br>delamination | n=28, N=5 | No | | |
| 3F | Frequency of change in sidedness | -60m to -40m<br>-40m to -20m | n=12, N=5<br>n=28, N=5 | No<br>No | Mann-Whitney test | 0.43118<br>(ns) |
| | | -40m to -20m<br>-20m to 0m | n=28, N=5<br>n=28, N=5 | No<br>Yes | | 0.00497<br>(*) |
| | | -60m to -40m<br>-20m to 0m | n=12, N=5<br>n=28, N=5 | No<br>Yes | | 0.00772<br>(*) |
| 3H | No. of NN/Total NN at start of constriction (%) | NN gained<br>by rearrangement<br><br>by division | <br>n=28, N=5<br><br>n=28, N=5 | <br>No<br><br>No | Mann-Whitney test | <br>0.01624<br>(*) |
| | | NN lost<br>as NB<br>by rearrangement | <br>n=28, N=5<br>n=28, N=5 | <br>No<br>Yes | | <br>1.17663E-7<br>(***) |
| | | NN gained<br>by rearrangement<br>NN lost<br>by rearrangement | <br>n=28, N=5<br><br>n=28, N=5 | No<br><br>Yes | | <br>3.1593E-7<br>(***) |
| 7I | % dpn$^+$ cells | Surface: Ctrl<br>mys$^{1mz}$ | N=10<br>N=10 | No<br>No | Mann-Whitney test | 5.65815E-4<br>(***) |
| | | Basal: Ctrl<br>mys$^{1mz}$ | N=10<br>N=10 | No<br>No | | 5.65815E-4<br>(***) |
| 7J | % pros$^+$ cells | Surface: Ctrl<br>mys$^{1mz}$ | N=10<br>N=10 | Yes<br>No | Mann-Whitney test | 0.01493<br>(*) |
| | | Basal: Ctrl<br>mys$^{1mz}$ | N=10<br>N=10 | Yes<br>Yes | | 7.68539E-4<br>(***) |
| 8H | % basal divisions | Ortho: Ctrl<br>mys$^{1mz}$ | N=10<br>N=10 | -<br>Yes | Two tailed Unpaired Student's t-test | (***) |
| | | Planar: Ctrl<br>mys$^{1mz}$ | N=10<br>N=10 | -<br>Yes | | (***) |
| 8I | % surface divisions | Ortho: Ctrl<br>mys$^{1mz}$ | N=10<br>N=10 | -<br>Yes | Two tailed Unpaired Student's t-test | (***) |
| | | Planar: Ctrl<br>mys$^{1mz}$ | N=10<br>N=10 | -<br>Yes | | (***) |
| 8L | Reversal of polarity | Surface: Ctrl<br>mys$^{1mz}$ | N=10<br>N=10 | -<br>No | Mann-Whitney test | 0.03484<br>(*) |
| | | Basal: Ctrl<br>mys$^{1mz}$ | N=10<br>N=10 | -<br>No | | 0.16808<br>(ns) |
| 8M | % symmetric divisions | Ctrl<br>mys$^{1mz}$ | N=10<br>N=10 | No<br>Yes | Mann-Whitney test | 0.01535<br>(*) |

Fig. 4C

| Mann-Whitney test | n=82<br>N=10 | n=100,<br>N=10 | n=41,<br>N=10 |
|---|---|---|---|
| | S4 | S5 | S6 |
| E vs D | 1.31E-04<br>(***) | 9.62E-04<br>(***) | 0.009<br>(**) |
| E vs C | 2.34E-04<br>(***) | 0.26705<br>(ns) | 0.3438<br>(ns) |
| E vs N | 6.06E-04 | 0.58425 | 1.53E-04 |

| | | | |
|---|---|---|---|
| | (***) | (ns) | (***) |
| D vs C | 0.03541 (*) | 0.00142 (**) | 0.13972 (ns) |
| D vs N | 0.11558 (ns) | 3.67E-04 (***) | 0.00619 (**) |
| C vs N | 0.84659 (ns) | 0.07576 (ns) | 0.00175 (**) |

| Mann-Whitney test | E | D | C |
|---|---|---|---|
| S4 vs S5 | 5.07E-04 (***) | 7.62E-04 (***) | 0.81431 (ns) |
| S5 vs S6 | 0.00204 (**) | 0.00239 (**) | 0.03268 (*) |
| S4 vs S6 | 0.02825 (*) | 0.01251 (*) | 0.0372 (*) |

| Morphometric State | Deadpan Heterogeneity | Normal Distribution |
|---|---|---|
| S4 | E | Yes |
| | D | No |
| | C | Yes |
| | N | No |
| S5 | E | No |
| | D | No |
| | C | No |
| | N | No |
| S6 | E | Yes |
| | D | Yes |
| | C | Yes |
| | N | No |

### Figure S1

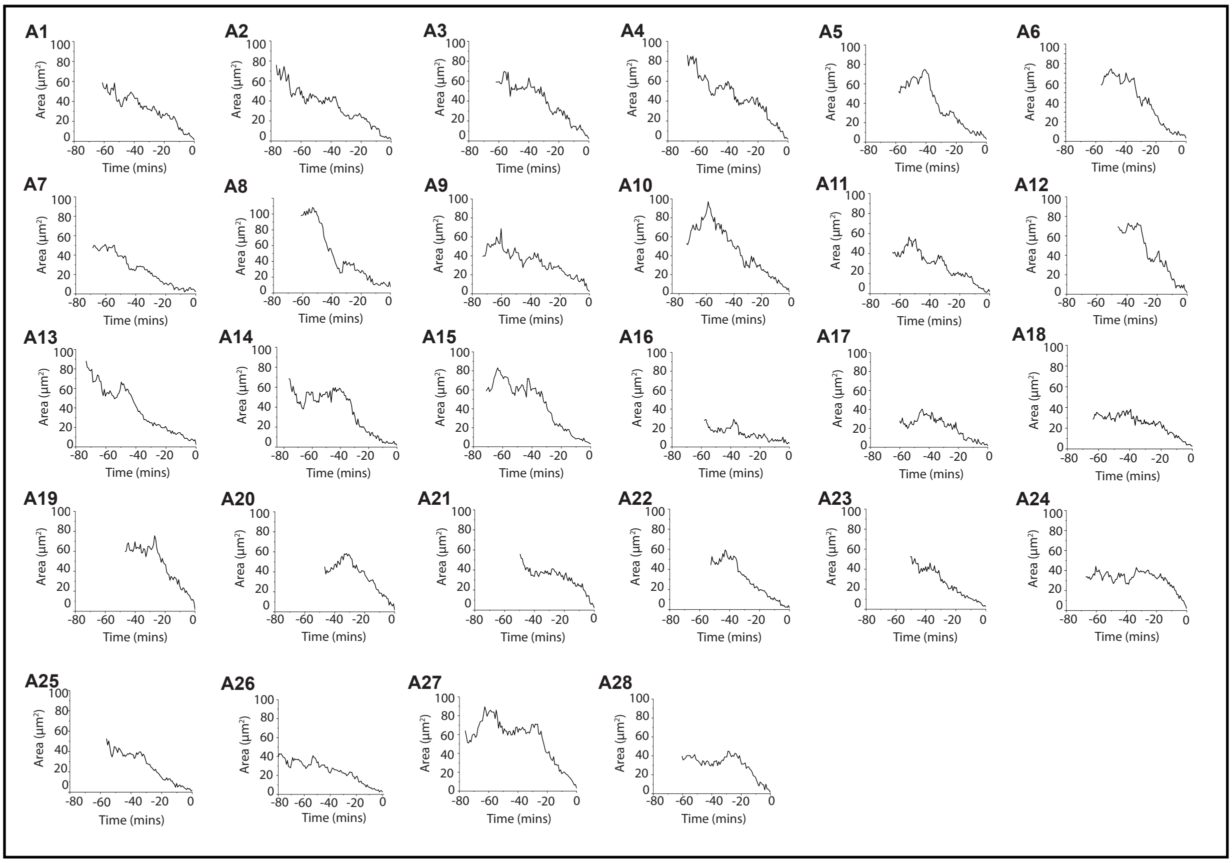

**Supplementary Figure 1: Apical area dynamics of delaminating neuroblasts** (accompanies Fig.2)

Apical area dynamics of delaminating neuroblasts from 5 embryos. A1-A15 show three constriction rates and A16-A28, a single constriction rate.

### Figure S2

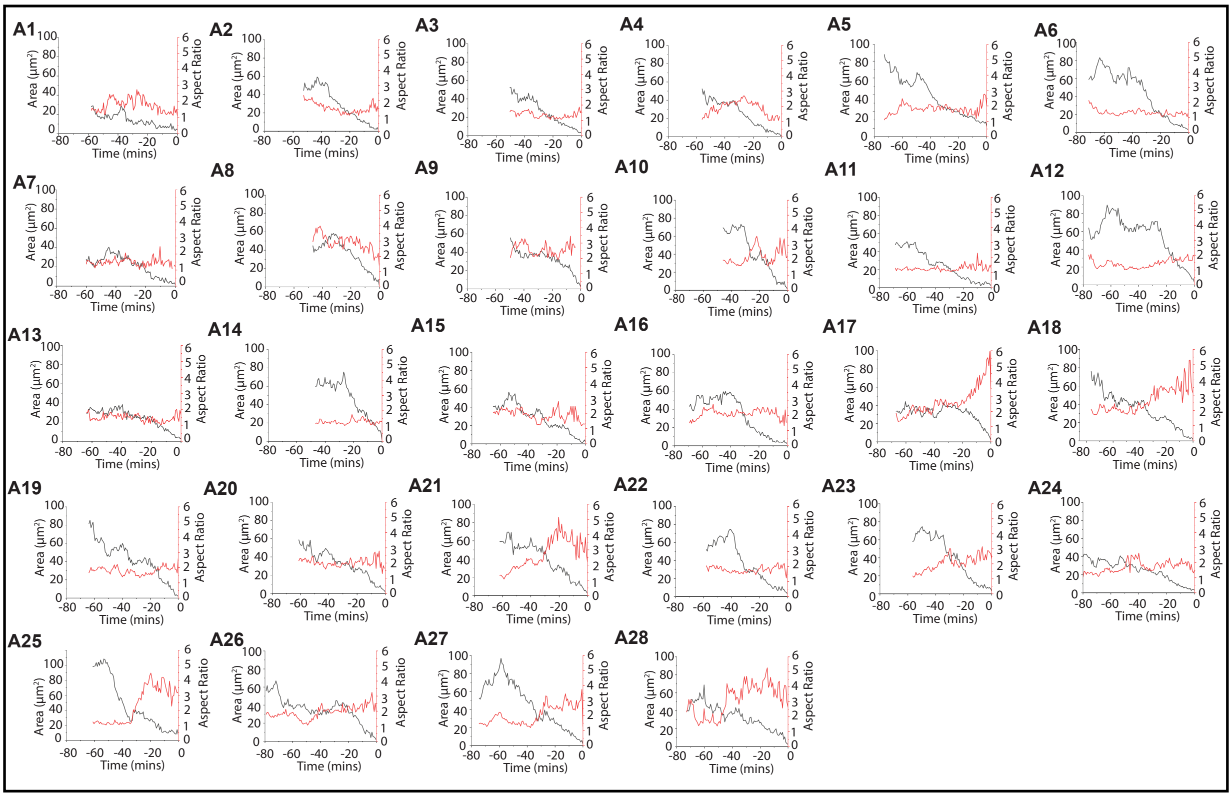

**Supplementary Figure 2: Apical area and aspect ratio dynamics in delaminating neuroblasts** (accompanies Fig.2)

(A1-A28) Quantitative morphodynamic analysis of 28 delaminating NBs from 5 embryos showing aspect ratio (red) and area (black) dynamics. t0 marks the time at which the area of the delaminating cell is less than 3 μm2. A1-A6, A7-A12 and A13-A28 show respectively a decrease, no change, or increase in shape anisotropy/aspect ratio in the 10 minutes prior to the end of delamination.

### Figure S3

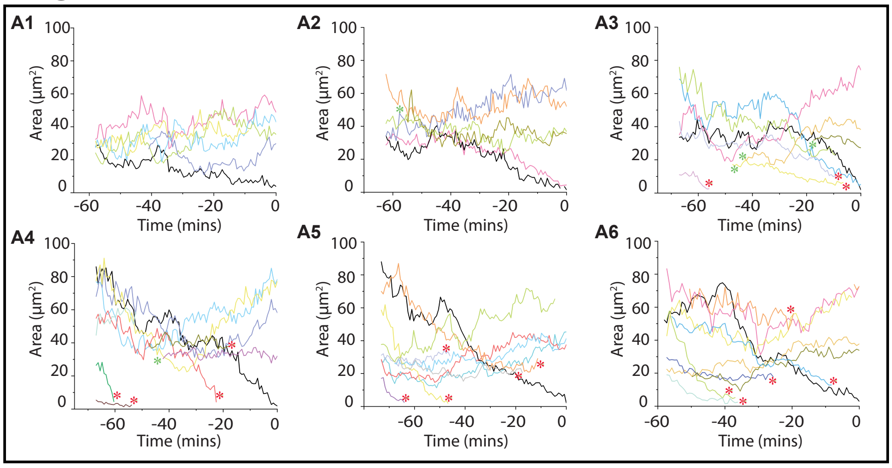

**Supplementary Figure 3: Apical area dynamics in NB cohorts** (accompanies Fig.3)

(A1-A6) Apical area dynamics of 6 delaminating NB cohorts. Black lines indicate the delaminating NB and the coloured lines are its nearest neighbours. Some nearest neighbours show an increase in area closer to the end of delamination. Asterisks mark neighbours lost.
